# Differential responses of transplanted stem cells to the diseased environment unveiled by a single molecular NIR II cell tracker

**DOI:** 10.1101/2020.03.12.988295

**Authors:** Hao Chen, Huaxiao Yang, Chen Zhang, Si Chen, Xin Zhao, Mark Zhu, Zhiming Wang, Xun Zhang, Yuebing Wang, Hung-Ta Wo, Kai Li, Zhen Cheng

## Abstract

Stem cell therapy holds high promises in regenerative medicine. The major challenge of clinical translation is to precisely and quantitatively evaluate the *in vivo* cell distribution, migration, and engraftment, which cannot be easily achieved by current techniques. To address this issue, for the first time, we have developed a single molecular cell tracker with a strong fluorescence signal in the second near-infrared (NIR-II) window (1000-1700 nm) for real-time monitoring of *in vivo* cell behaviors in both healthy and diseased animal models. The NIR-II tracker (CelTrac1000) has shown complete cell labeling with low cytotoxicity and profound long-term tracking ability for 30 days in high temporospatial resolution for semi-quantification of the biodistribution of primary mesenchymal stem cell and induced pluripotent stem cell-derived endothelial cells. Taking advantage of the unique merits of CelTrac1000, the responses of transplanted stem cells to different diseased environments have been discriminated and unveiled. Furthermore, we also demonstrate CelTrac1000 as a universal and effective technique for ultrafast real-time tracking of the cellular migration and distribution in a single cell cluster resolution, along with the lung contraction and heart beating. As such, this single molecular NIR-II tracker will shift the optical cell tracking into a single cell cluster and millisecond temporospatial resolution for better evaluating and understanding stem cell therapy, affording optimal doses and efficacy.

**Significance Statement:** For the first time, we synthesized a NIR-II tracker (CelTrac1000) for ultrafast real-time tracking of the migration trajectory of transplanted mesenchymal stem cells in the circulatory system with a single cell cluster resolution. Taking advantage of the merits of CelTrac1000, the responses of transplanted stem cells to different diseased environments, including acute lung injury, myocardial infarction, and middle cerebral artery occlusion, have been discriminated and unveiled in mice models. As such, our approach can help correlate critical biomedical information in stem cell therapies, such as stem cell dosing and engraftment and their relationships with efficacy, providing more accurate therapeutic treatment and outcomes in certain diseases during a long evaluation period (>30 days) in comparison with the commercial Qtracker (7-10 days).

## Introduction

Stem cells are pluripotent cells with self-renewing capacity, which can differentiate into a range of cell types under defined circumstances.^1-3^ In practice, stem cell therapy is promising for treating numerous disorders (*e.g.*, aging, autoimmune, and inherited disease) because autologous or allogeneic transplants can overcome the limitations of immune incompatibility.^4-9^ To date, stem cell therapy has been intensively employed in the treatment of wounds, blood and cardiovascular diseases, cartilage defect, diabetes, *et al*. Recently, stem cell therapy was on track for approval for human use in Japan for damaged corneas^10,11,12-17^. However, one roadblock preventing further widespread applications of stem cell therapy is the difficulty in tracking cell fates upon transplantation, preventing early assessment of dosage, retention, and therapeutic efficacy. Addressing this issue would greatly benefit the prognosis and overall outcomes.^18^ Hence, it is of great significance to track stem cells both *in vitro* and *in vivo* with high temporal and spatial resolution in a real-time manner during a prolonged period,, which can unveil the responses and behaviors of transplanted cells upon exposure to various diseased environments.

Noninvasive cell tracking is indispensable in stem cell therapy, including direct and indirect labeling techniques.^19^ By utilizing magnetic resonance imaging (MRI), single-photon emission computed tomography imaging (SPECT), positron emission tomography-computed tomography (PET-CT), and optical imaging, direct labeling strategies possess the advantages of abundant cell trackers and minimal interference with cells.^20, 21^ In practice, each cell tracking approach has its unique strengths and weakness. For instance, superparamagnetic iron oxides (SPIOs) as MRI trackers can ensure excellent anatomic information in deep organs, but the signal becomes ambiguous when cell numbers are low.^22, 23^ PET/CT detection is ultrasensitive, they are potential ionizing hazards and the temporal resolution is relatively low.^24, 25^ On the other hand, fluorescence imaging enjoys the merits of high sensitivity, high temporal resolution, and excellent maneuverability, which promotes their broad applications in a variety of biomedical imaging tasks in animal models. To realize *in vivo*, noninvasive, deep tissue fluorescence imaging with high temporal-spatial resolution, exogenous probes with emission in near-infrared (NIR) region are preferred.^26^ Specifically, great attention has been attracted to explore fluorescence probes in the second near-infrared (NIR-II) region (1000-1700 nm), which shows further improved tissue penetration depth and signal-to-noise ratio.^27^ Recently, we have demonstrated the first-in-human NIR-II fluorescence imaging-guided liver tumor surgery using organic single molecular indocyanine green (ICG), showing its advantages over traditional NIR-I imaging in clinical applications.^28^ But to date, most reported NIR-II probes are based on inorganic or organic nanomaterials: carbon nanotubes, quantum dots, and organic nanoparticles. However, this raises concerns over toxicity and surface modification complexity in *in vivo* imaging applications.29, 30 NIR-II quantum dots (Ag2S, PbS) for stem cell labeling have shown fine tracking results. But their unknown long-term toxicity brings substantial concerns due to the uncertain excretion of those heavy metal based inorganic nanoparticles.^31, 32^ In comparison, organic single molecular probes have well-defined components, high purity, clear excretion pathways and low cytotoxicity, which facilitate their applications in translational research. Certain organic, NIR-II emissive molecules with good biocompatibility have been reported for vascular structure imaging, showing great potential in *in vivo* applications.^33, 34^ Advancing the use of organic NIR-II emissive molecules as cell trackers into the field of stem cell therapy would significantly aid the development of new stem cell therapies. However, to the best of our knowledge, no organic NIR-II fluorescent trackers have yet been reported for *in vivo* stem cell tracking in literature.

Here, we have designed a novel NIR-II probe, CelTrac1000, for tracking stem cells in the animal models. CelTrac1000 is composed of a human serum albumin (HSA) molecule incorporated with a small molecule NIR-II dye (CH-4T) and further derivatized with a Tat peptide (**Fig. 1*A***). The structure of the single molecular tracker, CelTrac1000 can be precisely controlled and its synthesis can be easily scaled up. The toxicity, stability, and labeling efficiency of CelTrac1000 were first evaluated in cell models, showing that it could efficiently label stem cells within a few hours and stay in the cytoplasm for up to 30 days, with minimal leaking and perturbation to cell functions. Importantly, high resolution *in vivo* fluorescence imaging revealed the migration trajectory of administrated cells in the mouse circulation system with a single cell cluster resolution, which has never been achieved before. Furthermore, animal models of acute lung injury (ALI), myocardial infarction (MI) and middle cerebral artery occlusion (MCAO) were created. Direct imaging and comparisons of the transplanted stem cell distribution in the healthy and diseased models were successfully demonstrated and evaluated in high resolution and sensitivity, unveiling the differential responses of transplanted stem cells to diseased environments. CelTrac1000 could greatly benefit preclinical and clinical translation, providing a novel biotechnique of ultrafast, long-term stem cell tracking as a breakthrough in this field.

**Figure 1.**
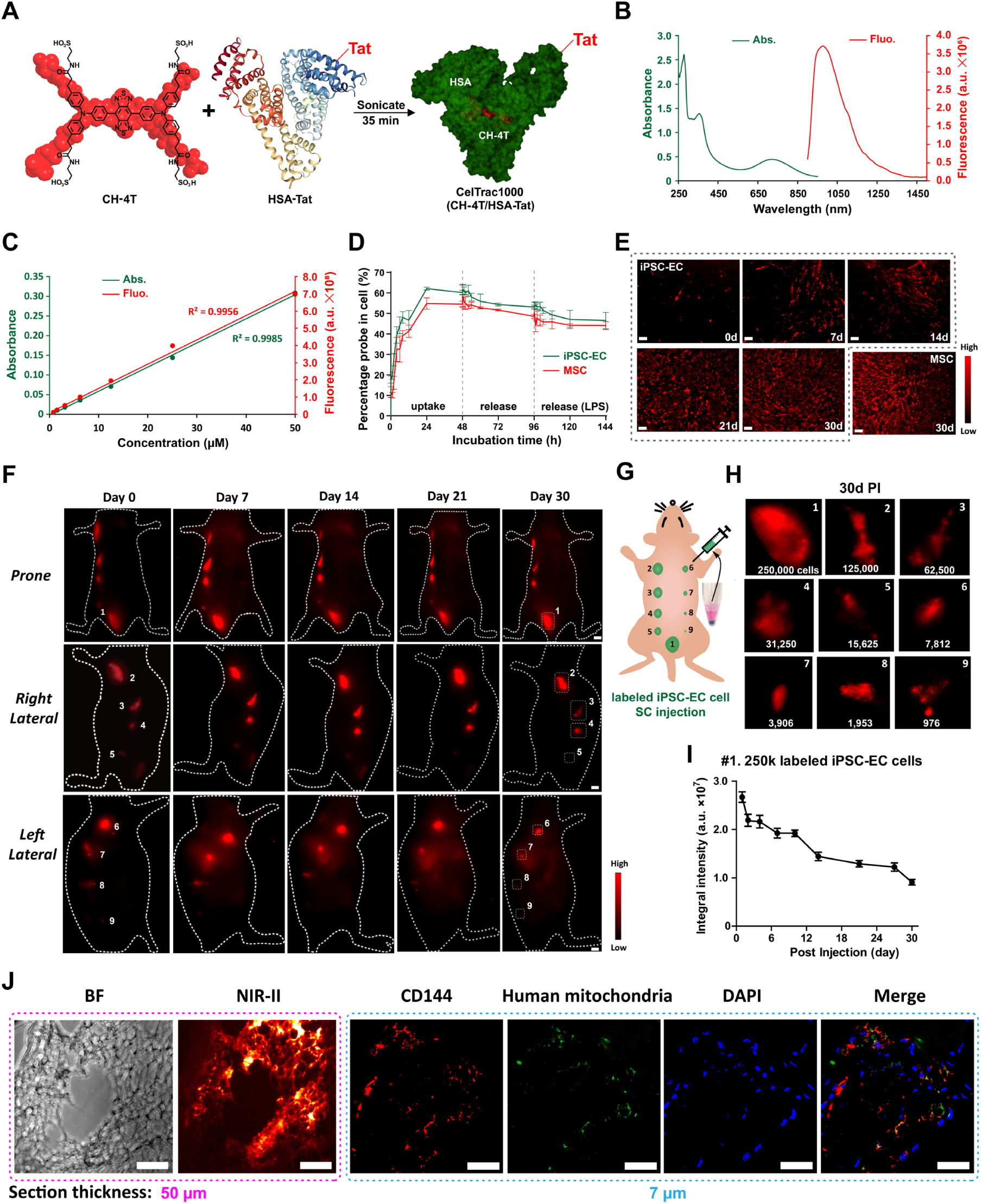
Synthesis of CelTrac1000 for long-term real-time cell tracking. (***A***) Chemical structure of CH-4T and preparation of the single molecule CelTrac1000 through integration with Tat-conjugated HSA. (***B***) Normalized UV-vis absorption and fluorescence spectra of CelTrac1000 in PBS (50 µM). (***C***) Plot of the maximum absorption and fluorescence intensities of CelTrac1000 at six different concentrations in PBS, indicating its linear response to concentration in a range of 0 to 50 µM. (***D***) Plot of the percentage of CelTrac1000 molecules in MSCs and iPSC-ECs under different treatments over 144h. The cells were first incubated with 50 µM of CelTrac1000 in culture medium for 48h, followed by incubation with probe-free culture medium until 96h. LPS (10 µg/mL) was then added into the culture medium to continue the release test until 144 h. (***E***) NIR-II fluorescent microscope images (785 nm excitation, 1000LP, 1000 ms) of CelTrac1000-labelled iPSC-ECs cultured for 0, 7, 14, 21, 30 days and the labeled MSCs after 30 days. (***F, G***) NIR-II fluorescent images (1000LP, 50 ms) and schematic illustration of CelTrac1000-labeled iPSC-ECs post subcutaneous injection at nine spots in a nude mouse. The cell numbers are 250,000, 125,000, 62,500, 31,250, 15,625, 7,812, 3,906, 1,953, and 976 at spots 1-9 accordingly. (***H***) Magnified NIR-II fluorescent images (1000LP, 100 ms) of the nine spots 30 days post injection. (***I***) Plot of the integral fluorescent intensities at spot 1 (250,000 iPSC-ECs) up to 30 days post injection. (***J***) Bright-field and NIR-II fluorescent microscope images (785 nm excitation, 1000LP, 1000 ms) of a skin section from spot 1 after 30 days post injection. The histological analysis of the skin section evaluated with immunofluorescent imaging of the ECs (CD144, red), injected iPSC-ECs (human mitochondria, green), and nuclei (DAPI, blue).

## Results

### Synthesis and characterization of CelTrac1000

The CelTrac1000 was synthesized through the incorporation of a NIR-II dye with a biocompatible protein carrier (**Fig. 1*A***). In brief, the NIR-II dye, CH-4T, was synthesized according to our previous report.^35^ Human serum albumin (HSA) was conjugated with Tat peptide (RKKRRQRRRC) through a carbodiimide-mediated coupling reaction to obtain Tat-HSA. The two components, CH-4T and Tat-HSA, were then mixed at equivalent concentrations and sonicated for 30 minutes (mins) in 1× PBS buffer at room temperature to afford CelTrac1000. The probe was analyzed by matrix-assisted laser desorption/ionization-time of flight mass spectrometry (MALDI-TOF-MS), suggesting it is a single molecular probe with a molar ratio of CH-4T : HSA : Tat at 1 : 1 : 1 (**Figs. S1-3** in Supplementary Information). We further examined the optical properties of CelTrac1000 in aqueous solution, showing absorption and emission peaks at 750 and 1000 nm, respectively (**Fig. 1*B***). Additionally, the absorbance and emission intensities of CelTrac1000 solutions showed a linear response to increased CelTrac1000 concentrations in a range of 0 to 50 µM (**Fig. 1*C***). This linear relationship between fluorescence and concentration can be attributed to the protection of HSA, which can significantly reduce the severe aggregation-caused quenching effect of fluorescent molecules. This unique optical signature is key to fluorescence semi-quantitative analysis, which is very challenging in practice.

An ideal cell tracker requires both high labeling efficiency and low cytotoxicity at working concentrations to ensure reliable results for precise analysis. We evaluated the direct labeling efficiency and toxicity of CelTrac1000 on human induced pluripotent stem cell-derived endothelial cells (iPSC-ECs) and mouse mesenchymal stem cells (MSCs) to assess optimized feeding concentrations. When the feeding concentration of CelTrac1000 increased from 0.78 to 200 µM, enhanced uptake efficiencies were observed in both of iPSC-ECs and MSCs within 48 h (**Fig. S4a**). On the other hand, the cell viability slightly decreased in both iPSC-ECs and MSCs when the concentration was above 100 µM (**Fig. S4b**). To avoid this, 50 µM was settled on as the optimal feeding dose for the following stem cell labeling. In addition, the CelTrac1000 showed negligible shifts in the gene expression profiles of the endothelial markers of iPSC-ECs (CDH5, PECAM, NOS3, KDR, NRG1, and ICAM1) and biomarkers of MSCs (CD44, ENG, LY6A/SCA-1) after labeling for two weeks (**Fig. S5**).

### *In vitro* cell tracking

First, the fluorescence stability of CelTrac1000 was evaluated in the biological environment to mimic long-term *in vivo* cell tracking. After incubation in 1× PBS buffer at 37 °C for two months, CelTrac1000 still showed excellent fluorescence intensity with negligible changes in emission profiles (**Fig. S6**). The uptake efficiency of CelTrac1000 in both iPSC-ECs and MSCs suggests that more than 40% of the probes (50 µM) were rapidly internalized into cells within the first 12 h (**Fig. 1*D***). At 48 h post incubation, the internalization percentage of CelTrac1000 in ECs and MSCs was 60% and 55%, respectively. In addition, approximately 6-7% of the probe was released from the cells into the fresh culture medium in the following 48 h. Overall, CelTrac1000 has shown excellent cell uptake and retention ability.

Customized NIR-II fluorescence microscopy was then used to study the performance of CelTrac1000 *in vitro*, tracking cells by recording fluorescence images of the labeled cells for up to 30 days post incubation. iPSC-ECs emitted intensive fluorescence with ∼100% labeling efficiency after overnight incubation with CelTrac1000 (day 0 in **Fig. 1*E***). Although the average fluorescence intensity from each EC gradually decreased due to cell proliferation in the next 30 days, almost all cells showed distinguishable fluorescence signals during the test period due to the excellent fluorescence stability and intracellular retention ability. The labeled MSCs also demonstrated a similar pattern of fluorescence changes in the *in vitro* culture, indicating a consistent performance of CelTrac1000 in the long-term labeling and tracking of different cell types. This is a significant advantage compared to commercial CellTracker and Qtracker, whose fluorescent signals only can last for a shorter period (∼7 days) according to previously reported *in vitro* results.^36-38^ As a result, the low toxicity, long-term tracking ability, and minimal leakage of CelTrac1000 presented it as a promising candidate for the next step of precise *in vivo* cell tracking.

### *In vivo* EC tracking and evaluation

A series of *in vivo* studies were designed to assess the performance of CelTrac1000 in the long-term tracking of animal models. First, different numbers of CelTrac1000-labeled iPSC-ECs (250000, 125000, 62500, 31250, 15625, 7812, 3906, 1953, 976) were subcutaneously injected at 9 spots on the backs of nude mice (**Figs. 1*F, G***). Intense fluorescent signals from the injection spots can be clearly distinguished from day 0 to day 30 post-transplantation (**Fig. 1*F***). Of note is all the injection sites emitted strong fluorescence on day 30 under a high magnification lens (**Fig. 1*H***), confirming the ultra-sensitive and stable imaging abilities of NIR-II CelTrac1000 in animal models. The fluorescence intensity changes in this study were further semi-quantitatively analyzed through the integration of the fluorescence intensities. As shown in **Fig. 1*I***, the integrated fluorescence intensity of spot one injected with 250,000 labeled ECs gradually decreased from day 0 to day 30, which was consistent with the previous studies. Analysis of spots 5 (15625 cells) and 9 (976 cells) as representative examples also demonstrated similar results (**Fig. S7**). The skin tissues at injection sites were collected at day 30 and the transplanted cell clusters were observed under customized NIR-II fluorescence microscope (**Fig. 1*J***). In addition, immunofluorescence staining of the tissue section suggests that a number of cells were double-positive with the expression of human mitochondria (green) and endothelial cell marker CD144 (**Fig. 1*J***), indicating that a certain number of human iPSC-ECs survived and engrafted during the test period. As such, labeling by CelTracker1000 did not affect the *in vivo* cell engraftment.

In the next step, a directed *in vivo* angiogenesis assay (DIVAA) was conducted to further visualize the *in vivo* angiogenesis of the iPSC-ECs. In brief, the semi-closed silicone angioreactors with a diameter of 5 mm were filled with a mixture of iPSC-EC and Matrigel, followed by subcutaneous implantation on both sides of the lower back of nude mice (**Fig. 2**). The survival of the iPSC-ECs and their participation in new vessel formation was longitudinally tracked at a depth of ∼ 4 mm for 30 days. Upon immunostaining of the iPSC-ECs in the transplanted angioreactors using isolectin-FITC, new vessel formation was observed from the iPSC-ECs 30 days post-surgery (**Fig. 2*H, I***). As such, our results validate the capability of transplanted iPSC-ECs to undergo angiogenesis as reported previously^39^ as well as provide the precise time window for angiogenesis, which can be a useful tool for better evaluating regenerative therapy with iPSC-ECs.

**Figure 2.**
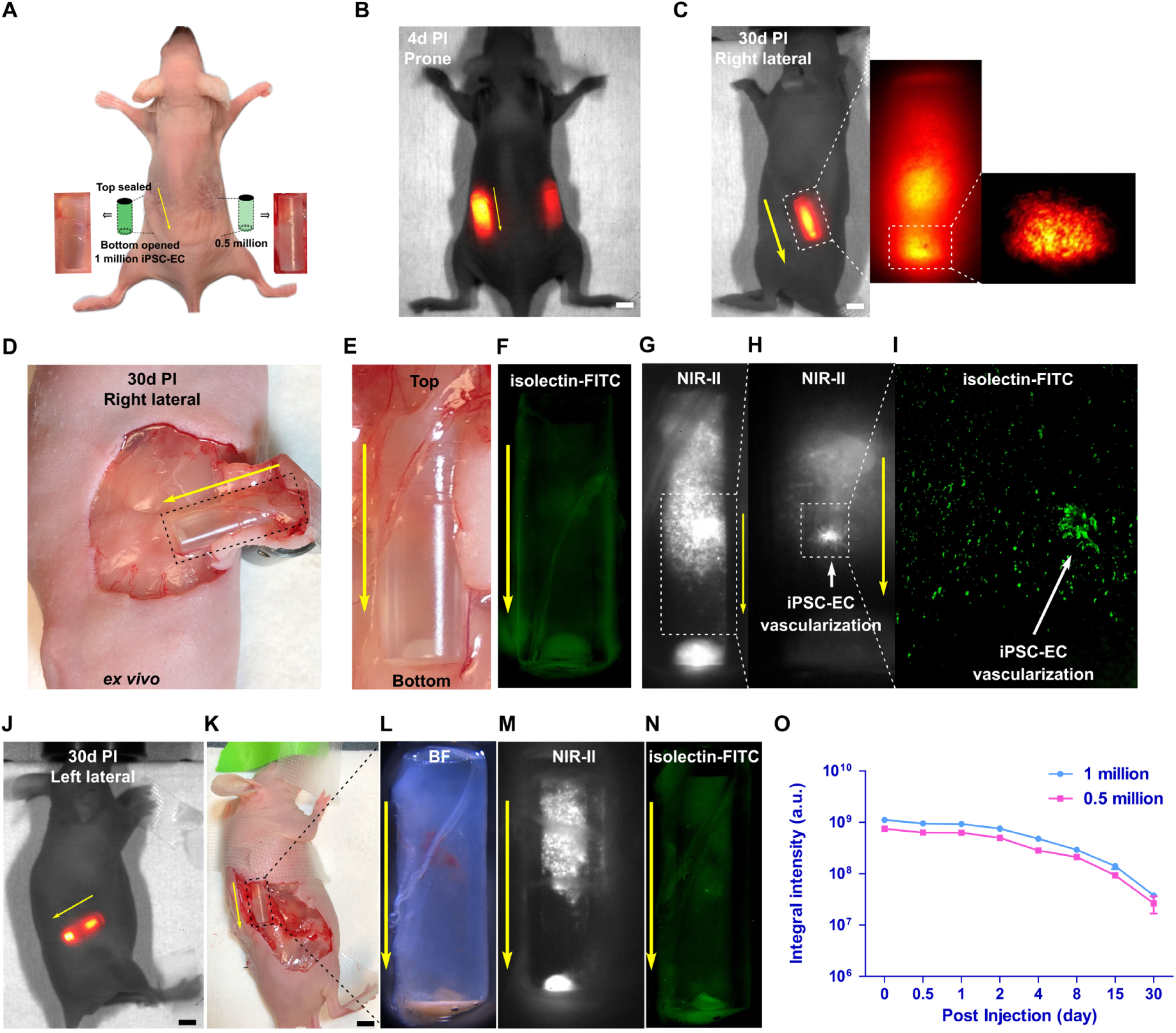
Directed *in vivo* angiogenesis assay for long-term evaluation of angiogenesis induced by iPSC-ECs. (***A***) A nude mouse implanted with two semi-closed silicone angioreactors in the left (1 million CelTrac1000-labeled iPSC-ECs) and right (0.5 million CelTrac100-labeled iPSC-ECs) lower back. (***B, C***) NIR-II fluorescent images (1000LP, 100 ms) of the same mouse 4 days and 30 days post angioreactor transplantation, respectively. The magnified NIR-II images show clear visualization of labeled iPSC-ECs in the reactor. (***D***) An immunofluorescent image of the iPSC-ECs after isolectin-FITC staining. (***E*-*H***) NIR-II fluorescent images (1000LP, 100 ms) of the extracted angioreactor from the left lower back, 30 days post transplantation. (***I***) Immunofluorescent microscopy image of iPSC-EC vascularization after isolectin-FITC staining. (***J*-*N***) NIR-II fluorescent images (1000LP, 100 ms) and immunofluorescent images of iPSC-EC after isolectin-FITC staining of the extracted angioreactor from the right lower back 30 days, post transplantation. (***O***) Plot of the integral NIR-II fluorescent intensities of the angioreactors (1 and 0.5 million labeled iPSC-ECs, n=4) up to 30 days post injection

### Highly sensitive and efficient *in vivo* MSC tracking in the circulation system

The whole body NIR-II fluorescence imaging was performed to track intravenously injected CelTrac1000-labeled MSCs to reveal the real-time cell migration in the circulatory system. 1,000,000 labeled MSCs were injected into the tail vein of each healthy mouse. The ultra-fast and sensitive NIR-II imaging systems allowed us to monitor the dynamic biodistribution of the administrated MSCs in real-time (**Fig. 3**). The lung showed strong fluorescence signals immediately after MSC injection as the majority of administrated cells were initially trapped inside lung capillaries.^40^ Later, the circulating MSCs gradually accumulated in the liver and spleen. More importantly, the real-time traffic of single-cell clusters at several positions in the circulation system was observed, as well as their migration through blood vessels from one organ to another (**Video S1** and **S2**). In particular, the migration of cells among the lungs, liver, and spleen, as well as the cell traffic within the hindlimb blood vessels can be clearly visualized (**Video S2**). **Figs. 3*A*** and **3*B*** show the representative frames of fluorescence images from Video S1 and S2, respectively. The single-cell clusters in high resolution are visualized (**Fig. 3*A***). Sequential frames clearly indicate the trajectory of three individual MSC clusters under magnification in the region highlighted in **Fig. 3*B***, suggesting the ultra-high sensitivity of our *in vivo* cell tracking technique. From the dimensions of the cell cluster (**Fig. 3*C***), it is estimated that one single cell cluster captured on the NIR-II images contained ∼1000 MSCs. However, considering the diffraction effect of light, one cell cluster observed under the imaging system could likely contain fewer cells. As a result, our approach can unveil *in vivo* cell migration behaviors in detail that would be impossible to be captured with conventional cell tracking techniques.

**Figure 3.**
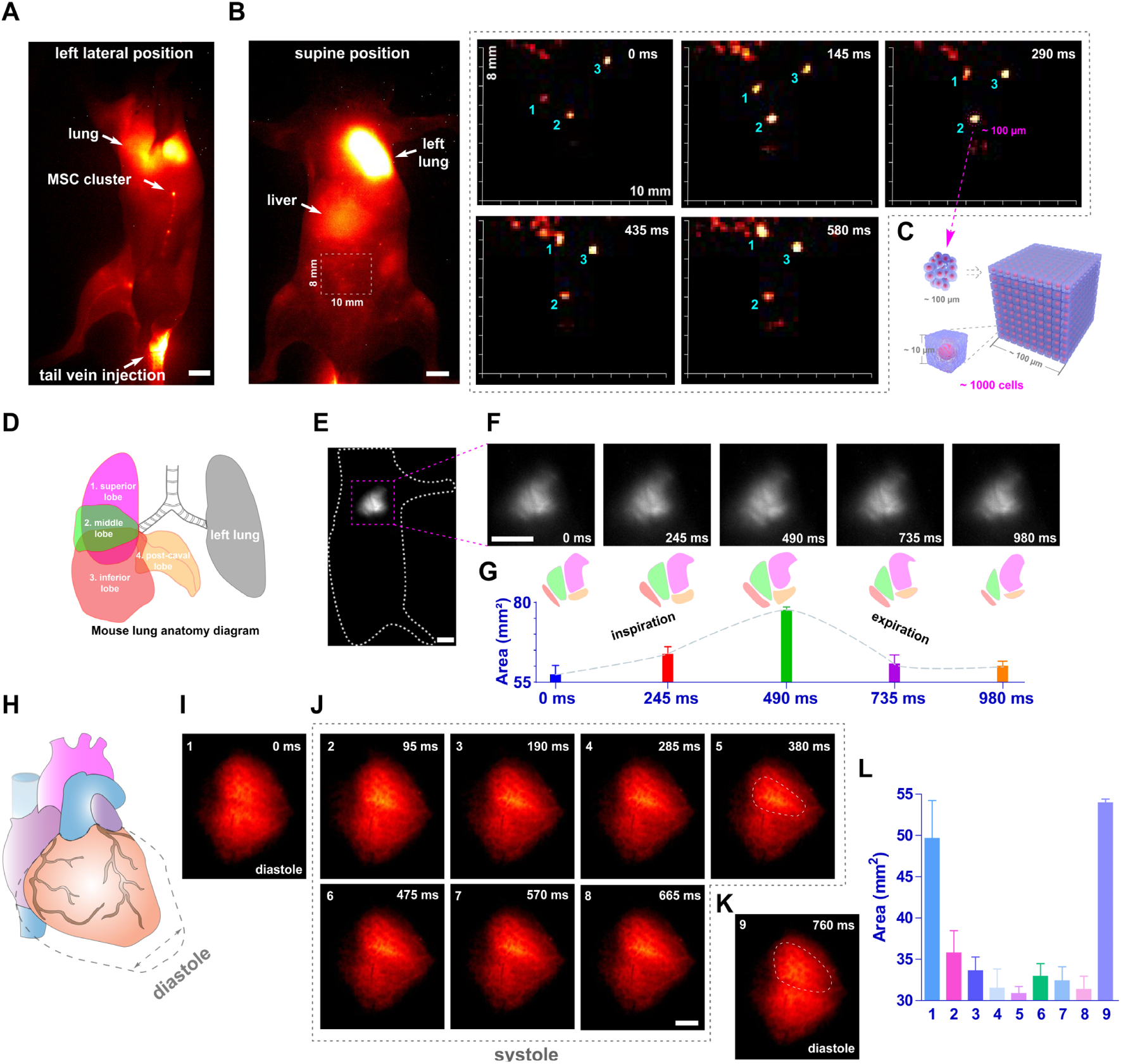
Dynamic stem cell tracking in mouse circulation system. (***A***) Representative whole-body NIR-II fluorescent image (1100LP, 100 ms) of a mouse 5 mins post intravenous injection of CelTrac1000-labeled MSCs, corresponding to the Supplementary Video S1. Scale bar: 5 mm. (***B***) The migration trajectory of MSC clusters in the circulatory system over 580 ms. The white outlined box (8 × 10 mm) is shown under high magnification. This corresponds to the Supplementary Video S2. (***C***) Schematic illustration of the magnified cell clusters. To facilitate the estimation of cell numbers, we assume that a single MSC is in a cubic shape with a side length of ∼10 µm. As the clusters shown in (***B***) suggest a dimension of ∼100 µm, we estimate the cell number in one cluster is ∼1000. (***D***) Schematic illustration of mouse lung anatomy. The right lung has 4 lobes: superior, middle, inferior and post-caval lobes. The left lung has 1 lobe. (***E***) Representative NIR-II fluorescent image (1100LP, 200 ms) of mouse lung 1.5h post intravenous injection of CelTrac1000-labeled MSCs. Scale bar: 5 mm. (***F***) The real-time monitoring of the lung lobe movement during inspiration and expiration over 980 ms, corresponding to Supplementary Video S3. Scale bar: 5 mm. (***G***) The trend of total lung lobe area changes during inspiration and expiration processes. Data were extracted from (***F***) for analysis. (***H***) Schematic illustration of systole and diastole phases of the cardiac cycle. (***I*-*K***) NIR-II fluorescent image (1000LP, 50 ms) of heart in diastole and systole phases over 760 ms, corresponding to Supplementary Video S4. Scale bar: 5 mm. (***L***) Quantitative analyses of the fluorescent areas of the heart in systole and diastole phases of the cardiac cycle, corresponding to ***I***-***K***.

Furthermore, the movement of the right lung lobes during inhalation and exhalation was recorded in real-time when the mouse was placed in a lateral position (**Figs. 3*D*-*F*, Video S3**). The schematic in **Fig. 3*D*** shows the structure of the right lung, consists of four lobes: the superior, middle, inferior, and post-caval lobes. The positions of four lobes relocate accordingly when breathing. Because of the high fluorescence intensity from the MSCs trapped in the lung, the movement of these lung lobes was captured under the ultra-fast NIR-II imaging system (**Fig. 3*F***). The high-resolution images allowed us to extract the profile of each lobe and calculate the size change during the inspiration/expiration cycle (**Fig. 3*G***). In addition, we placed the mouse in a supine position and focused on the heart to record the heart beating throughout the full cardiac cycles. Thanks to the presence of administrated MSCs in blood and their strong NIR-II fluorescent signal, it was clearly observed and recorded the diastole and systole phases of the heart under anesthesia status (**Figs. 3*H*-*K*, Video S4**). These results prove the highly efficient and accurate imaging of MSCs in the mouse circulation system, suggesting that our NIR-II cell tracking approach can serve as a simple and promising technique to reveal obfuscated biological mechanisms and processes.

### *In vivo* MSC tracking in disease models

One intrinsic quality of MSC is their ability to homing to injured sites, secreting a broad spectrum of paracrine factors to create a regenerative microenvironment.^41^ The CelTrac1000 was thus used to evaluate the biodistribution and retention of labeled MSCs in injured mice of various disease models, including acute lung injury (ALI), myocardial infarction (MI), and middle cerebral artery occlusion (MCAO) model of stroke. All these models were confirmed through hematoxylin and eosin (H&E) staining examination.

First, lipopolysaccharide (LPS) was used to create ALI in C57BL/6J mice through intratracheal instillation. Upon intravenous administration of the labeled MSCs (1 million) into ALI and healthy mice, NIR-II imaging studies were performed to monitor the dynamic changes of fluorescent signals in the animal bodies (**Fig. 4*A***). We then carried out quantitative analyses of the average fluorescent intensities in different organs using ImageJ, revealing the details of their dynamic cell migration and retention of the MSCs in the lung, heart, spleen, and liver (**Fig. 4*B***). *Ex vivo* images of organs and H&E staining of sectioned tissues were acquired after *in vivo* imaging study on day 3 (**Figs. 4*C*-*H***). As shown in **Fig. 4*B***, the lungs trapped a number of MSCs in both ALI and healthy mice in the first 5 mins post-injection. The fluorescence intensity from the injured lungs of ALI mice was significantly stronger compared to that from the healthy mice at 6 h post-injection. This can be mainly attributed to the homing capacity of MSCs to injured sites in the lung lobes, resulting in higher engraftment (**Figs. 4*C*-*D***). We were able to clearly observe the MSC clusters under the NIR-II imaging system in the lung tissue sections collected from the ALI mouse 72 h post cell administration (**Fig. 4*C***). On the contrary, more MSCs escaped from lungs in the healthy control and re-entered into the circulation system, resulting in a rapid decrease in fluorescence intensity of the healthy lungs from 3 h post cell administration (**Fig. 4*B***).

**Figure 4.**
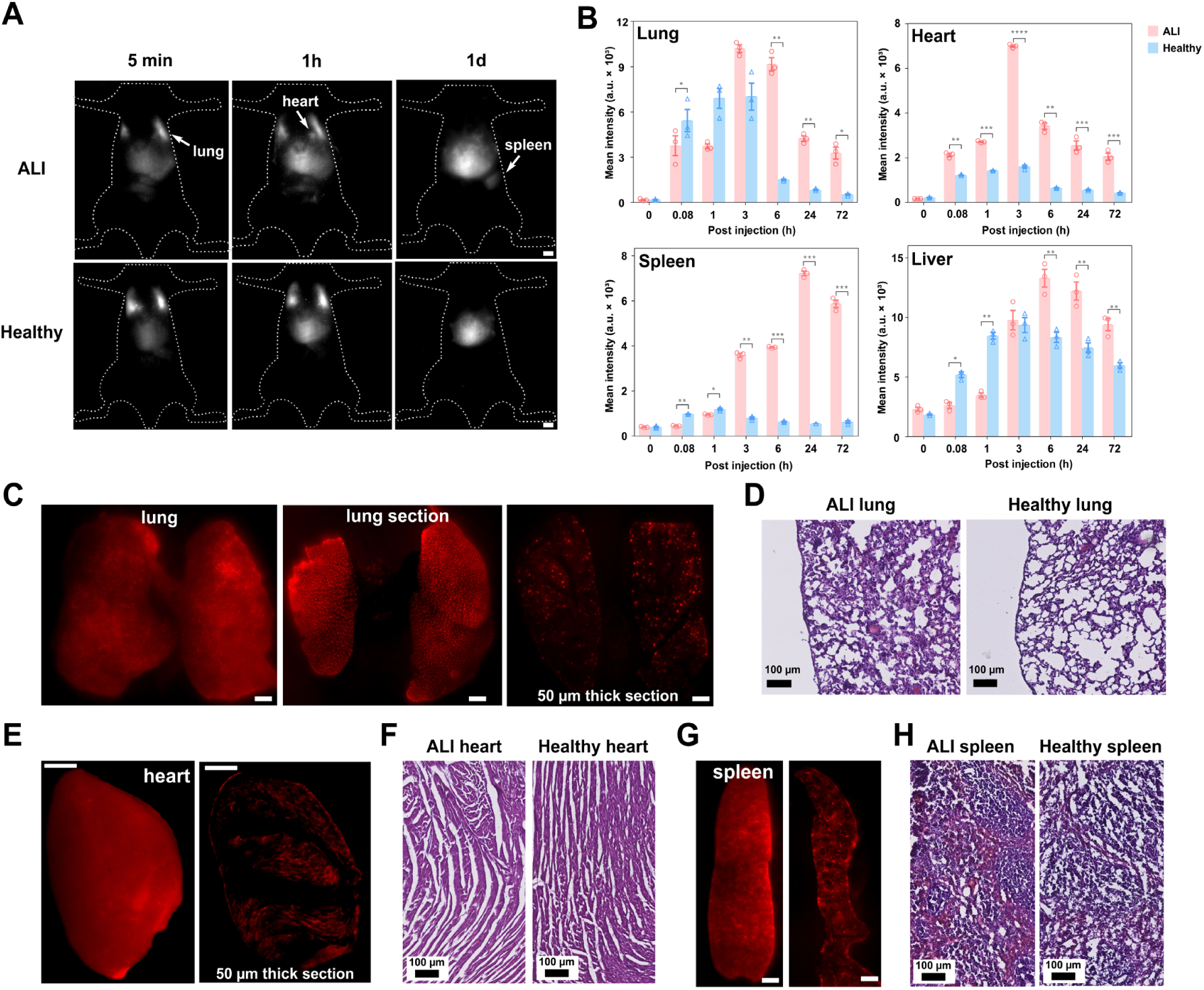
Unveiling the biodistribution of MSCs in the mouse model of acute lung injury (ALI). (***A***) Representative NIR-II fluorescent images (1000LP, 100 ms) of the ALI mouse and healthy mouse upon intravenous injection of CelTrac1000-labeled MSCs (1 million), showing the distinct biodistribution signatures. Scale bar: 5 mm. (***B***) Quantitative analyses of the average fluorescent intensities from the organs (lung, heart, spleen, liver) at different times intervals (5 min, 1 h, 3 h, 6 h, 24 h, and 72 h post cell injection in the ALI and healthy mice, respectively. n = 3, \**p* < 0.05, \*\**p* < 0.01, \*\*\**p* < 0.001, \*\*\*\**p* < 0.0001. (***C***) NIR-II fluorescent images (1000LP, 200 ms) of the whole lung, sectioned lung tissue (2 mm in thickness), and cryo-sectioned lung tissue (50 µm in thickness) from the ALI mouse. Scale bar: 5 mm. (***D***) H&E stained lung tissues from ALI and healthy mice. (***E***) NIR-II fluorescent images (1000LP, 200 ms) of the whole heart and cryo-sectioned heart tissue (50 µm in thickness) from the ALI mouse. Scale bar: 5 mm. (***F***) H&E stained heart tissues from ALI and healthy mice. (***G***) NIR-II fluorescent images (1000LP, 200 ms) of the whole spleen and cryo-sectioned spleen tissue (50 µm in thickness) from the ALI mouse. Scale bar: 5 mm. (***H***) H&E stained spleen tissues from ALI and healthy mice. The mice were sacrificed to collect organs 72h post injection of the MSCs for *ex vivo* analyses.

Surprisingly, the heart and spleen of ALI mice showed intense fluorescent signals after 3 h post-injection, with significant differences comparing to that of healthy mice. Looking at the heart, the fluorescent intensity of the heart from the ALI mice gradually increased until 3 h, then decreased thereafter. Of note is that the signals from the ALI heart were significantly stronger than those from the healthy heart at all time points post cell injection. These results could be due to the fact that cardiac dysfunction and damage was also caused in the LPS-induced ALI model, which prompted MSC migration and homing to the injured heart tissues^42^. Similar results were observed in the spleen tissues, with significantly higher fluorescent intensity from injured spleen in ALI mice. At 72 h post cell injection, significant cell accumulation was observed in the liver and spleen, which are the primary excretory organs. As a result, our findings provided direct evidence to confirm that ALI model creation using LPS caused inflammation in major organs that led to the recruitment of MSCs.

To further confirm if the homing of MSCs in heart tissues of ALI mice was caused by inflammation and injury, a MI mouse model was created for MSC transplantation. In a parallel experiment, we injected CelTrac1000, instead of labeled MSCs, into the MI mice for comparison. The biodistribution and dynamic behavior of the probe and MSCs in MI mice showed very distinct patterns (**Fig. 5*A***). Similar to what we discovered in the ALI model, the administrated labeled MSCs in MI mice were trapped in the lung and heart immediately upon intravenous injection while the injected probe only showed minimal intensity in the lung and heart 5 mins post-injection (**Figs. 5*B*** and **5*C***). The fluorescent signals from the hearts of MI mice gradually increased post CelTrac1000 injection. This could be caused by the high concentration of the probe in the blood, which when combined the higher blood vessel permeability of the injured heart tissues, may facilitate leakage of the probe, accumulating in the surrounding tissues. In addition, the relative fluorescence intensity ratio of heart to lung in the MI model was significantly higher than that of the ALI model 6 h post-MSC injection (p < 0.05) (**Fig. 4*B*, Fig. 5*B***, and **Fig. 5*C***), due to the severer heart injury created in the MI model. The fluorescence signal from the heart was still high at 72 h post-MSC injection, suggesting the MSCs were preferentially attracted to the infracted myocardium. This result was further confirmed from *ex vivo* fluorescence imaging and H&E staining results of heart tissue sections (**Figs. 5*E*** and **5*F***). Meanwhile, the liver’s signals remained at a high level from the initial injection up to 3 days in both CelTrac1000-injected and MSC-injected mice. Overall, the CelTrac1000 labeling strategy allowed us to monitor the dynamic migration and distribution of MSCs in a MI model, with high sensitivity and specificity, which has never been achieved by fluorescence imaging before.

**Figure 5.**
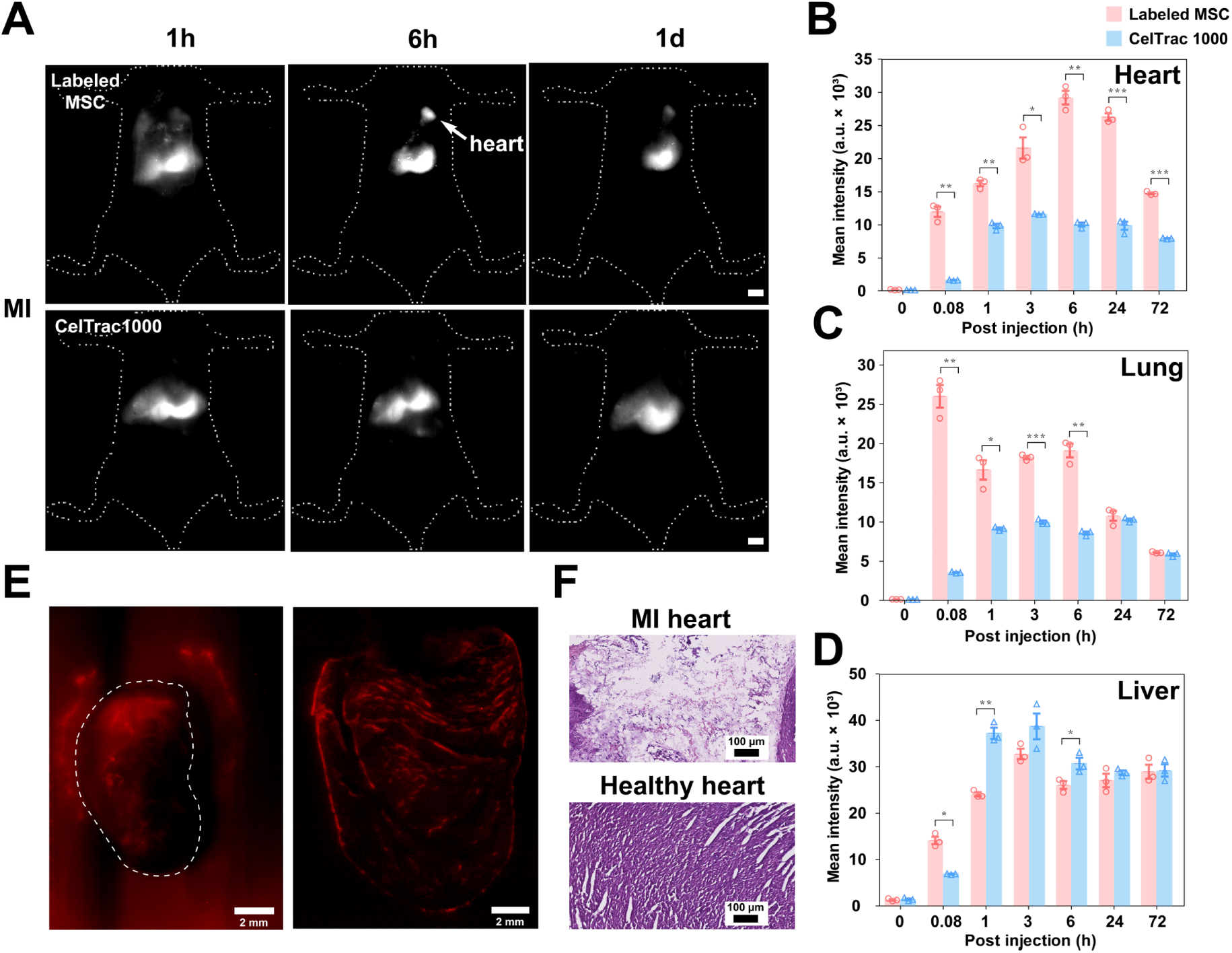
Unveiling the biodistribution of MSCs in the mouse model of myocardial infarction (MI). (***A***) Representative NIR-II fluorescent images (1000LP, 100 ms) of the MI mice upon intravenous injection of CelTrac1000-labeled MSCs (2 million) and CelTrac1000 probe (0.2 µmol), respectively. Scale bar: 5 mm. Quantitative analyses of the average fluorescent intensities from heart (***B***), lung (***C***), and liver (***D***) at different time intervals (5 min, 1h, 3h, 6h, 24h, and 72h) post injection of CelTrac1000-labeled MSCs and CelTrac1000 probes, respectively. n = 3, \**p* < 0.05, \*\**p* < 0.01, \*\*\**p* < 0.001, \*\*\*\**p* < 0.0001. (***E***) NIR-II fluorescent image (1000LP, 200 ms) of the whole heart and cryo-sectioned heart tissue (50 µm in thickness) from the MI mouse injected with labeled cells. Scale bar: 2 mm. (***F***) Images of H&E stained heart tissues from the MI and healthy mice for comparison. The mice were sacrificed to collect organs 72h post injection of the MSCs for *ex vivo* analyses.

Ischemic stroke is one of the major causes of mortality in developed countries and the leading cause of long-term disability worldwide.^43^ To investigate the *in vivo* cell migration and distribution in the treatment of ischemic stroke, we created a middle cerebral artery occlusion (MCAO) mouse model, creating a stroke in the left cerebral hemisphere, according to the established protocol^44^. The through-skull *in vivo* cerebrovascular fluorescence imaging of MCAO mice was carried out after intravenous administration of CelTrac1000-labeled MSCs. At 5-min post-MSC injection, the left hemisphere of the brain with the MCAO showed disrupted vascular structure, while the intact right hemisphere exhibited healthy vascular structure co-localized with the presence of labeled MSCs in the blood flow (**Fig. 6*A***). At 30-min post-MSC injection, apparent migration of the MSCs to the stroke site of the left cerebral hemisphere was discovered. The MSCs were distributed across the whole brain in both the left and right hemispheres 1 h post cell administration (**Fig. 6*B***). Cerebral ischemia is known to induce dramatic activation and release of various cytokines, chemokines and adhesion molecules.^45, 46^ The inflammatory mediators released in the ischemia area can modulate the permeability of the blood brain barrier (BBB).^47^ As a result, the dispersed MSCs across the whole-brain can be attributed to the enhanced BBB permeability and intense inflammatory reaction caused by the stroke.

**Figure 6.**
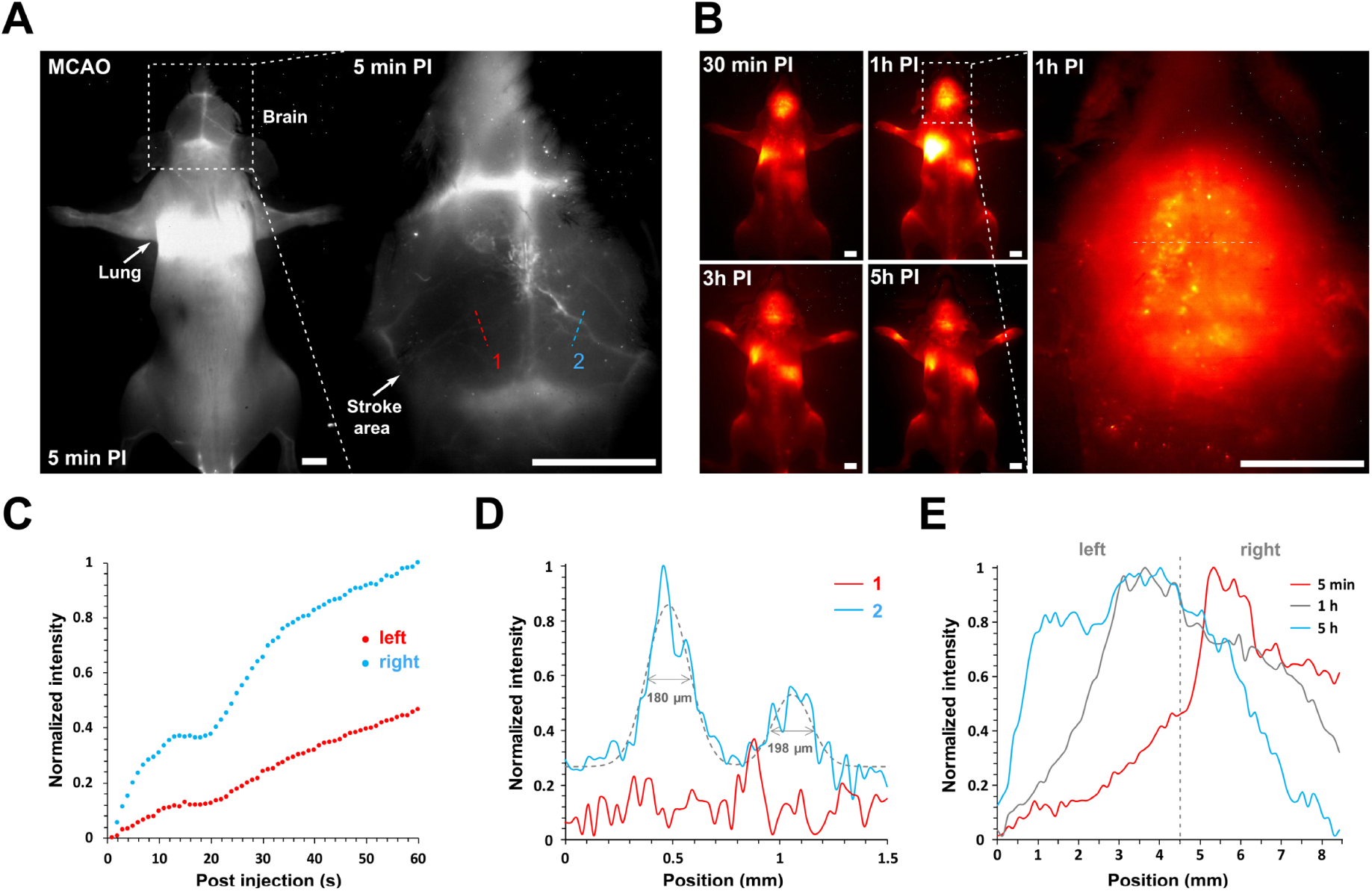
Unveiling the brain vascular structure of a MCAO mouse model and MSC response to brain inflammation in a MCAO mouse. (***A***) Representative NIR-II fluorescent images (1100LP, 200 ms) of a MCAO model mouse 5 min post intravenous injection of CelTrac1000-labeled MSCs (2 million). Note that the blood vessels in the left cerebral hemisphere of the stroke site are not visible while the right cerebral hemisphere can be clearly observed. Scale bar: 5 mm. (***B***) Representative NIR-II fluorescent images (1100LP, 200 ms) of the MCAO mouse at different time points (30 min, 1h, 3h, and 5h) post intravenous injection of CelTrac1000-labeled MSCs. Scale bar: 5 mm. (***C***) Time course of fluorescence intensity in the brain blood vessels of a MCAO mouse corresponding to lines 1 and 2 in ***A***. (***D***) The cross-sectional NIR-II fluorescent intensity profiles of locations 1 and 2 in ***A***. (***E***) Time course of fluorescence intensity in the brain of a MCAO mouse corresponding to the white dashed line in ***B***, revealing the dynamic intensity changes in the left and right hemispheres.

We further quantitatively evaluated the dynamic fluorescence intensity changes in the left and right hemispheres of the MCAO mice. The maxima of fluorescence cross-sectional intensities of the blood vessels from both hemispheres gradually increased in the first 60 s post-MSC injection due to the enrichment of the MSCs within brain vessels (**Fig. 6*C***). In addition, the fluorescent signal of blood vessels in the right hemisphere showed much higher intensities as compared to those in the left hemisphere because the blockage of vessels constrained the blood flow at the stroke site. The cross-sectional NIR-II fluorescent intensity profile of line 2 in **Fig. 6*A*** showed a full-width half-maximum (FWHM) of 180 and 198 µm (**Fig. 6*D***), indicating the ultrahigh imaging sensitivity of the labeled MSCs. Quantification of the cross-sectional NIR-II fluorescent intensity profile across the left and right hemispheres (the white line in the magnified brain image in **Fig. 6*E***) can give us more details of the dynamic process of MSC migration in the brain. It suggests that the fluorescence intensity in the right hemisphere was much higher than that in the left hemisphere 5 min post-MSC injection, due to the higher concentration of MSCs in normal blood vessels compared to the blocked ones. Over time, the fluorescent intensity in the right hemisphere gradually decreased while the left hemisphere increased at 5 h post-injection. This phenomenon can be caused by the gradually decreasing concentration of MSCs in normal blood vessels, while the homing of MSCs at the stroke site became more and more significant due to enhanced BBB permeability and the triggered inflammation.

As a result, our approach provides a feasible and ultra-sensitive technique to reveal the details of MSC migration and distribution in different disease models, which are hardly observed with conventional imaging methods. This opens a new avenue to facilitate and assist clinicians in providing a more precise treatment and outcome assessment when applied in stem cell therapy.

## Discussion

In summary, a facile stem cell labeling and *in vivo* tracking technique has been realized to unveil the homing and migration of stem cells upon transplantation in mice models of various diseases. The single molecular tracker scales well for translational research and applications with simple synthesis and easily controlled quality. CelTrac1000 has shown its robust cell labeling and tracking capability on both stem cell-derived endothelial cells and primary mesenchymal stem cells, with low cellular toxicity and minimal effects on cell functions. Taking advantage of the merits of NIR-II fluorescence imaging, we successfully demonstrate that the single molecular NIR-II tracker can visualize the migration trajectory of single-cell clusters in the circulatory system with high sensitivity and temporal/spatial resolution. More importantly, the differential MSC distributions and migrations have been imaged, analyzed and compared in both healthy and ALI, MI, and MCAO mice models in detail. This can help correlate critical biomedical information, such as stem cell dosing and engraftment and their relationships with efficacy, in stem cell therapies, providing more accurate therapeutic treatment and outcomes in certain diseases. Overall, our strategy provides researchers and clinicians a promising stem cell tracking approach to promote the translational potential of stem cell-based therapies in the near future.

## Materials and Methods

### Synthesis and characterization of CelTrac1000

The water-soluble organic NIR-II dye, CH-4T, was synthesized as reported^35^. HSA was conjugated with a Tat peptide through a two-step EDC/NHS coupling reaction in 10× PBS buffer. In brief, Tat peptide (RKKRRQRRRC trifluoroacetic acid salt from GenicBio, China) (43.4 mg) was dissolved in 0.5 mL of dry DMSO with 89.2mg EDC and 7.14 mg NHS for 1.5 h at room temperature. The activated Tat peptide was then mixed with 100mg HSA (dissolved in 2 mL of 10× PBS). After reacting overnight at room temperature, the obtained Tat-HSA molecules were purified with an Amicon Ultra-5 mL 10k to eliminate the excess free Tat peptides, EDC and NHS. To prepare 0.2 mL of 1mM CelTrac1000 solutions, the obtained Tat-HSA (13.6 mg) and CH-4T (0.28 mg) were mixed in 1× PBS buffer (0.2 mL). The mixture was sealed in a 1.5 mL sterilized centrifuge tube and put into a bath sonicator for 30 min to afford CelTrac1000. The stock solution was then stored at 4 °C for further use.

### Characterization of CelTrac1000

The molecular weights of HSA, Tat-HSA, and CelTrac1000 (Tat-HSA/CH-4T) were measured by matrix-assisted laser desorption/ionization time of flight mass spectrometry (MALDI-TOF MS). The absorption spectrum of CelTrac1000 in water was recorded on an ultraviolet-visible-NIR Cary 6000i spectrophotometer. The NIR-II fluorescence spectrum was recorded on a customized spectroscope, with excitation at 808 nm and a power output of ∼160 mW. The excitation laser was filtered with a combination of an 850/1,000/1,100/1,200/1,300/1,350/1,400 nm long-pass filters (Thorlabs). The sample was loaded into a 1 cm path-length cuvette and the signal was filtered through a 910 nm long-pass filter (Thorlabs) to reject the incident excitation laser light. The emitted signal was recorded on a spectrometer (IsoPlane SCT-320) coupled to a liquid nitrogen-cooled InGaAs detector array (Princeton Instruments, NIRvana TE 640). Upon acquisition of the raw data, a correction file was applied to correct the variable InGaAs quantum efficiency as a function of detection wavelength, as well as the variable 910 nm long-pass filter extinction features across the NIR-II spectral region. A series of absorption and emission profiles of CelTrac1000 solutions were measured at probe concentrations of 0, 1.5625, 3.125, 6.25, 12.5, 25, and 50 µM.

### NIR-II imaging

Mice tail vein was infused with a venous catheter for IV injection of labeled cells or probes. All NIR-II images were collected on a 640 × 512 pixels two-dimensional InGaAs array (Princeton Instruments, NIRvana TE 640). The excitation laser was an 808 nm laser diode at a power density of ∼140 mW cm^-2^. Emission was typically collected with 1000, 1100 nm LP filter (Thorlabs). A prime lens (50 mm or 75 mm, Edmund Optics) was used to obtain magnifications ranging from 1× (whole-body) to 2.5× (high magnification) magnification by changing the relative position of the camera, lens, and animals. A binning of 1 and the variable exposure time was used for the InGaAs camera (640 × 512 pixels) to capture images in the NIR-II window. Images were processed with ImageJ (NIH).

### NIR-II fluorescent microscopy imaging

A Nikon ECLIPSE Ni fluorescent microscope with an InGaAs camera (Princeton Instruments, NIRvana TE 640), 785 nm, 100 mW/cm^2^ laser excitation, 1000 nm long-pass filter (Thorlabs), 800 nm short-pass filter (Thorlabs) and 805 nm cut-on long pass dichroic mirror (Thorlabs) was used for NIR-II fluorescent microscopy imaging.

### Study of cellular uptake and leakage of CelTrac1000

Mouse mesenchymal stem cells (MSCs) and human-induced pluripotent stem cell-derived endothelial cells (iPSC-ECs) were seeded in 6-well plates individually at a density of 0.5 million cells per well (n = 4 each). When the cells reached 80% confluence, 50 µM of CelTrac1000 in 2 mL of culture medium was added into each well. At designated time intervals (0, 0.5, 1, 2, 4, 6, 8, 12, 24, and 48 h), medium (100 µL) was collected from each well for further analysis of uptake efficiency through fluorescence measurement. At 48 h, each well was washed with PBS buffer, and 3 mL of fresh culture medium was added. Upon addition of the fresh medium, medium (100 µL) was then collected from each well at 0, 0.5, 1, 2, 4, 6, 12, 24, and 48 h for further analysis of leakage from cells. After washing, lipopolysaccharide (LPS) was diluted in 3 mL of medium (10 µg/mL) and added into each well for culture. After 0, 0.5, 1, 2, 4, 6 h, medium (100 µL) was collected from each well. The LPS medium was then discarded and 3 mL of fresh medium was added into each well, followed by a collection of 100 µL of the medium at 12, 24, and 48 h. The collected samples were analyzed to obtain fluorescence intensities for analysis of probe leakage from cells after treatment with LPS. The calculation formulas are in the supporting information.

### Cytotoxicity of CelTrac1000

The metabolic activity of MSCs and ECs was evaluated by CellTiter 96^®^ AQueous One Solution Cell Proliferation Assay (Promega) individually. MSCs and ECs were seeded in 96-well plates at 2 × 10^4^ cells/mL. After 24 h incubation, the medium was replaced by CelTrac1000 solution at concentrations of 200, 100, 50, 25, 12.5, 6.25, 3.125, 1.5625 and 0.78125 µM, and the cells were then incubated for 48 h. The cells were washed twice by 1× PBS buffer, followed by addition of a mixture of 20 µL CellTiter 96^®^ Aqueous reagent and 100 µL culture medium into each well. After further incubation for 1.5 h, the absorbance was recorded by a microplate reader at 490 nm (n = 6 in each loading concentration). The cell viability was expressed as the ratio of absorbance from the cells incubated with CelTrac1000 to that of the cells incubated with culture medium only.

### Effect of CelTrac1000 on gene expression of iPSC-EC and MSC

The cellular responses of CelTrac1000 treatment on the transcriptional levels of iPSC-ECs were measured by the real-time quantitative PCR (RT-qPCR). First, RNA was extracted using the RNeasy Mini Kit (Qiagen). All the RNAs used in this study were A260/280 = 1.9 ∼ 2.1. Then 500 ng cDNA was synthesized via reverse transcription using the iScript cDNA Synthesis Kit (Bio-Rad). The qPCR was performed with the TaqMan gene expression assay, and the mRNA expression levels of CDH5, KDR, PECAM, NRG1, NOS3, and ICAM1 were examined. For MSCs, the mRNA expression levels of CD29, CD44, CD105, Sca-1 were analyzed. The final results were demonstrated as the relative expressions to the control group (N = 3 in each group).

### Long-term cell tracking

MSCs and ECs were individually cultured in 6-well plates to achieve 80% confluence. After medium removal and washing with 1× PBS buffer, 50 µM of CelTrac1000 in culture medium was then added to the wells. After overnight incubation at 37 °C, the cell monolayers were washed twice with 1× PBS buffer and cultured in fresh medium for 7, 14, 21, and 30 days, respectively. After designated time intervals, the NIR-II fluorescence images of cells were recorded upon excitation at 785 nm with a 1000 nm long-pass filter.

To further investigate the detection limit of labelled cells in *in vivo* studies, different amounts of CelTrac1000-labelled ECs (250,000, 125,000, 62,500, 31,250, 15,625, 7,812, 3,906, 1,953, 976) were subcutaneously injected on the back of nude mice (n=4). The fluorescence images of these 9 spots were then recorded at designated time intervals (0, 7, 14, 21, 30 days) by a customized NIR-II small animal imaging facility upon excitation at 808 nm with a 1000 nm long-pass filter. The skin tissues were collected on day 30 for histological analysis.

### Animal handling

All animal experiments were approved by Stanford University’s Administrative Panel on Laboratory Animal Care. Eight-week-old female C57BL/6 mice and BALB/c nude mice were purchased from Charles River for imaging studies and housed at the Research Animal Facility of Stanford University.

### *In vivo* angiogenesis assay

The directed *in vivo* angiogenesis assay (DIVAA) was conducted to visualize the process of angiogenesis in nude mice (n=4). iPSC-ECs were first labeled by 50 µM of CelTrac1000 in culture medium at 37 °C for 12 h, followed by trypsinization to collect the cell suspensions. A semi-closed silicone strength (sealed top and open bottom, 5 mm in diameter) was filled with labeled iPSC-ECs and Matrigel. Two silicone angioreactors were separately loaded with different cell numbers (0.5 million and 1 million). The angioreactors were then subcutaneously implanted at both sides of the lower back of nude mice. The mice were imaged under a customized NIR-II small animal imaging facility upon excitation at 808 nm with an 1100 nm long-pass filter at designated time intervals. After 1 and 7 months, the mice were sacrificed to collect the angioreactors for *ex vivo* evaluation of the regenerative effect of transplanted iPSC-ECs.

### Surgical procedures of animal models

Female C57BL/6 mice (n=4 each group) were used to create acute lung injury (ALI), myocardial infarction (MI), and middle cerebral artery occlusion (MCAO) models for imaging studies, following the procedures reported in the literature^48-50^. In brief, ALI was induced by giving LPS in 1× PBS solution through intratracheal instillation at a dose of 2 mg/kg. The LPS-treated mice were housed for another 24 h before imaging studies. The sub-acute MI model was achieved *via* coronary ligation for 60 minutes, followed by reperfusion. In the surgery, a polypropylene suture was passed from the left fringe of the pulmonary infundibulum to the lower right of the left auricle for ligation. The mice were then housed for 4 days before stem cell therapy and imaging. The silicon-tripped intraluminal thread occlusion method was employed to establish the MCAO mouse model. An 11 mm silicone-coated nylon thread was introduced into the left common carotid artery of the mouse and directed into the internal carotid artery until it obstructed blood flow to the middle cerebral artery. After 60 minutes, the filament was withdrawn and wounds were sutured. The mice were then housed for another 24 h before imaging studies. All the surgical procedures were performed when the mice were under anesthesia. H&E staining of organs was performed to confirm the success of disease models after imaging studies.

### *In vivo* dynamic tracking of MSCs in the circulation system of mice

Before imaging studies, the hair of C57BL/6 mice was shaved using a depilatory gel. During imaging processes, the mice (n=4) were placed on an imaging stage connected with an electric heating pad to maintain a consistent temperature. Mouse MSCs isolated from C57BL/6 mice were expanded and incubated with 50 µM of CelTrac1000 in culture medium at 37 °C for 12 h, followed by trypsinization to collect the cell suspensions. The labeled MSCs (2 million) were suspended in 1× PBS (200 µL) and injected into the healthy mouse through the tail vein. Real-time NIR-II fluorescent images were recorded by a customized NIR-II small animal imaging facility upon excitation at 808 nm with a 1000 nm or 1100 nm long-pass filter with an exposure time of 50 ms, 100 ms or 200 ms. The real-time monitoring of the injected MSCs in blood vessels, movement of lung lobes during inspiration and expiration, and heartbeat behavior were investigated to reveal the dynamic behavior of MSCs in the circulation system.

### *In vivo* cell tracking in disease models

MSCs were first incubated with 50 µM of CelTrac1000 in culture medium at 37 °C for 12 h, followed by trypsinization to collect the cell suspensions. The labeled MSCs (1 million) were suspended in 1× PBS (200 µL) and injected into the ALI/MI/MCAO mouse model through the tail vein. The fluorescent images of mice were then recorded at designated time intervals (0, 5 minutes, 1 h, 3 h, 6 h, 1 day and 3 days) by a customized NIR-II small animal imaging facility upon excitation at 808 nm with a 1000 nm long-pass filter (ALI/MI) or 1100 nm long-pass filter (MCAO). In the ALI model study, the same amount of labeled MSCs was intravenously injected into each healthy mouse for biodistribution comparison.

### Histological analyses of engraftment of iPSC-ECs

The engraftment of iPSC-ECs in nude mice (n=4) skin tissues was investigated to evaluate their participation in angiogenesis. The mice were sacrificed at day 30 post iPSC-EC injection. The full-thickness skin tissues from the cell injection spots were collected and placed in an Optimal Cutting Temperature compound (Thermo Fisher Scientific, Hampton, NH, USA) on dry ice for embedding and freezing. The blocks were then cryo-sectioned into sections at 10 µm thickness for immunofluorescence staining processes. The slides were fixed in an ice-cold acetone/methanol mixture (50%/50%) and stained with primary anti-mouse CD144 antibody (MAB9381, 1:100, R&D Systems) and secondary donkey-anti-mouse Alexa Fluor^®^ 594 (A21203, 1:200, Thermo Fisher Scientific), anti-human mitochondria antibody Alexa Fluor^®^ 488 conjugate (MAB1273A4, 1:100, EMD Millipore), and DAPI.

### Statistical Analyses

Data from different groups were analyzed by the student’s *t*-test, and differences at the 95% confidence level (*p* < 0.05) were considered to be statistically significant.

## Acknowledgments

This publication was supported by the National Natural Science Foundation of China (31870991, 81301160), Shanghai Pujiang Program (19PJ1411100), American Heart Association (AHA) Postdoctoral Fellowship Award (18POST34030106), Stanford University, Department of Radiology. We also would like to thank Dr. Andrew Olsen from Stanford Neuroscience Microscopy Service (NIH NS069375) on the support of confocal imaging of iPSC-ECs.

## Supporting Information

**Figure S1.**
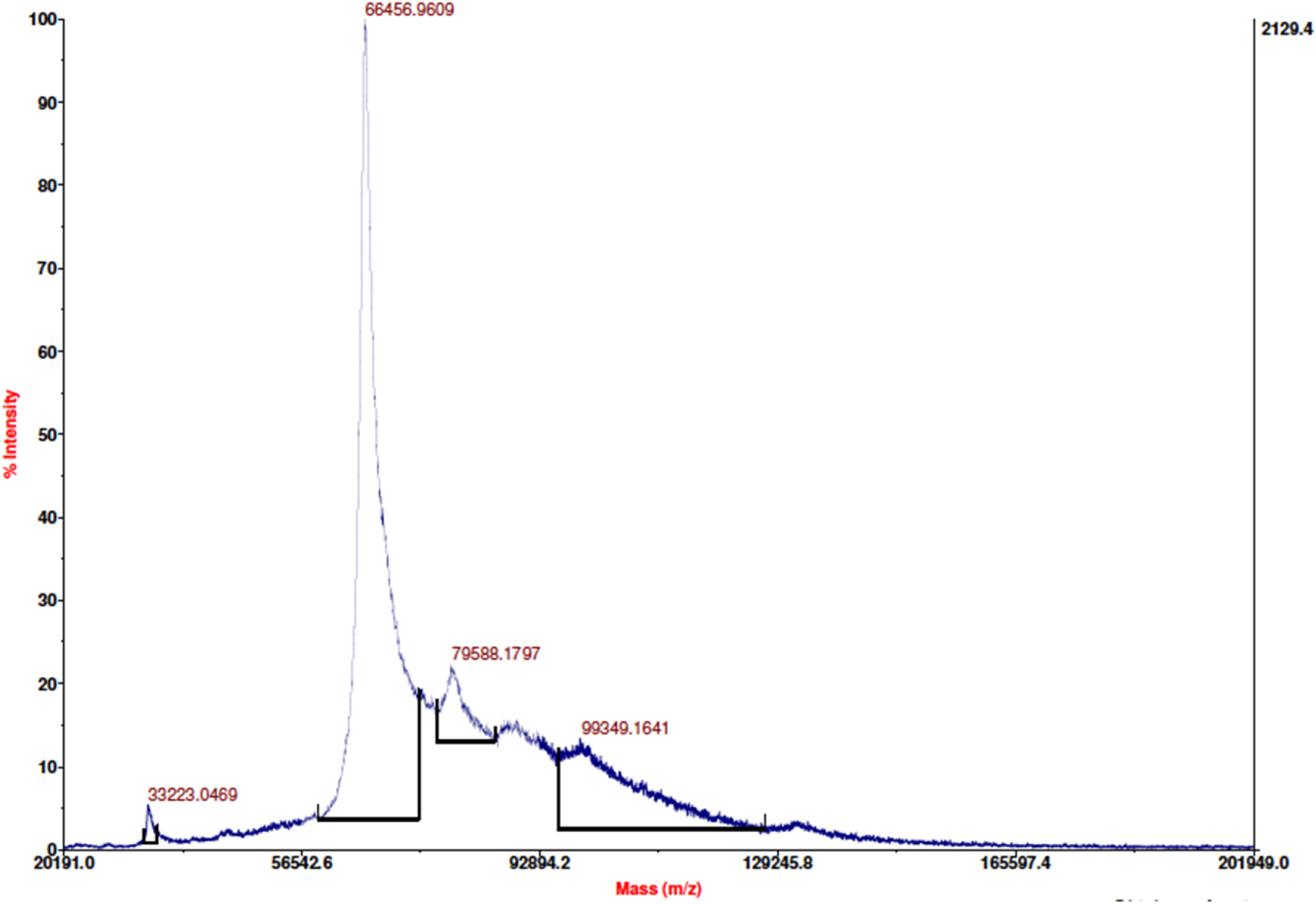
MALDI-TOF-MS spectrum of HSA.

**Figure S2.**
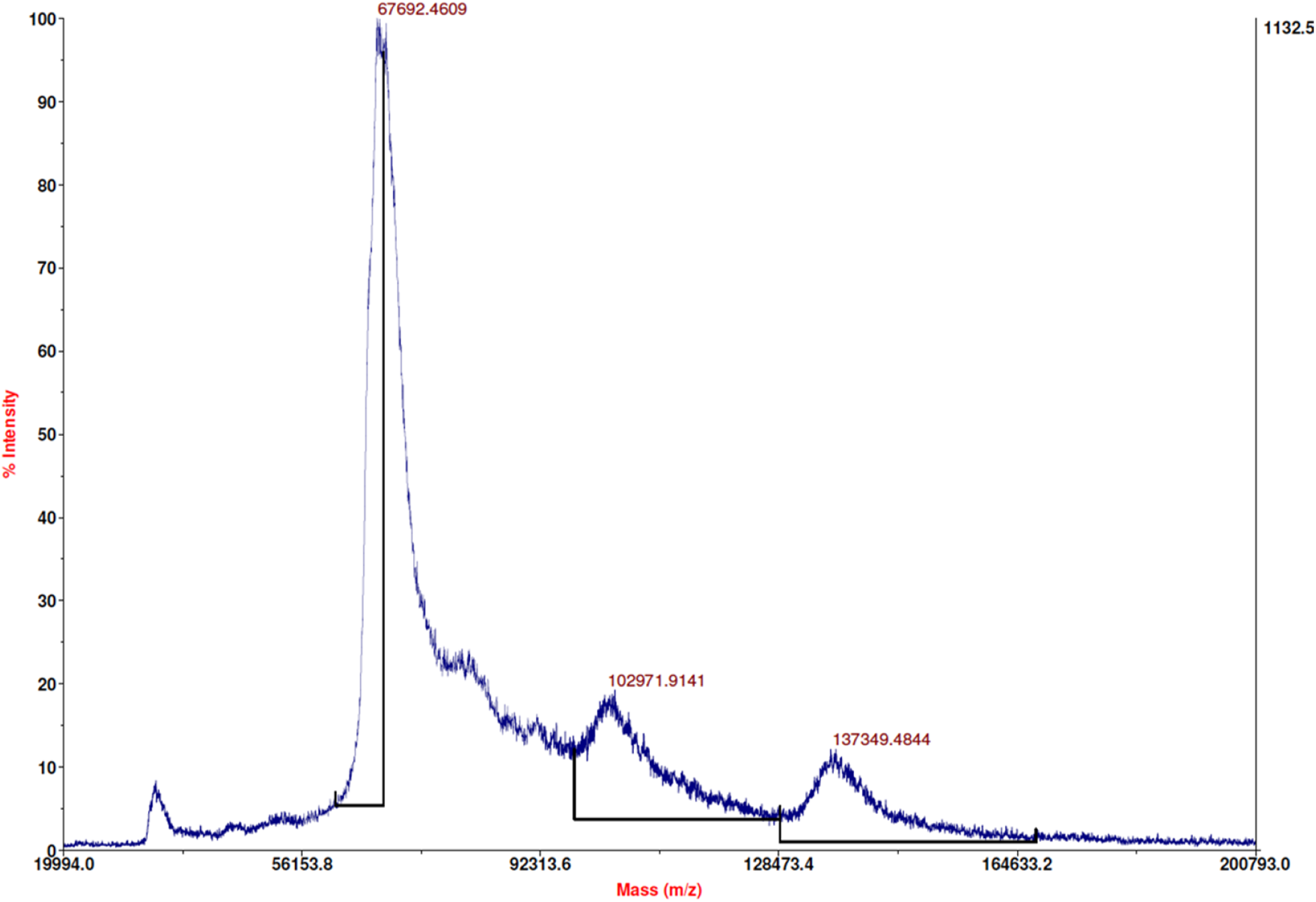
MALDI-TOF-MS spectrum of HSA-Tat.

**Figure S3.**
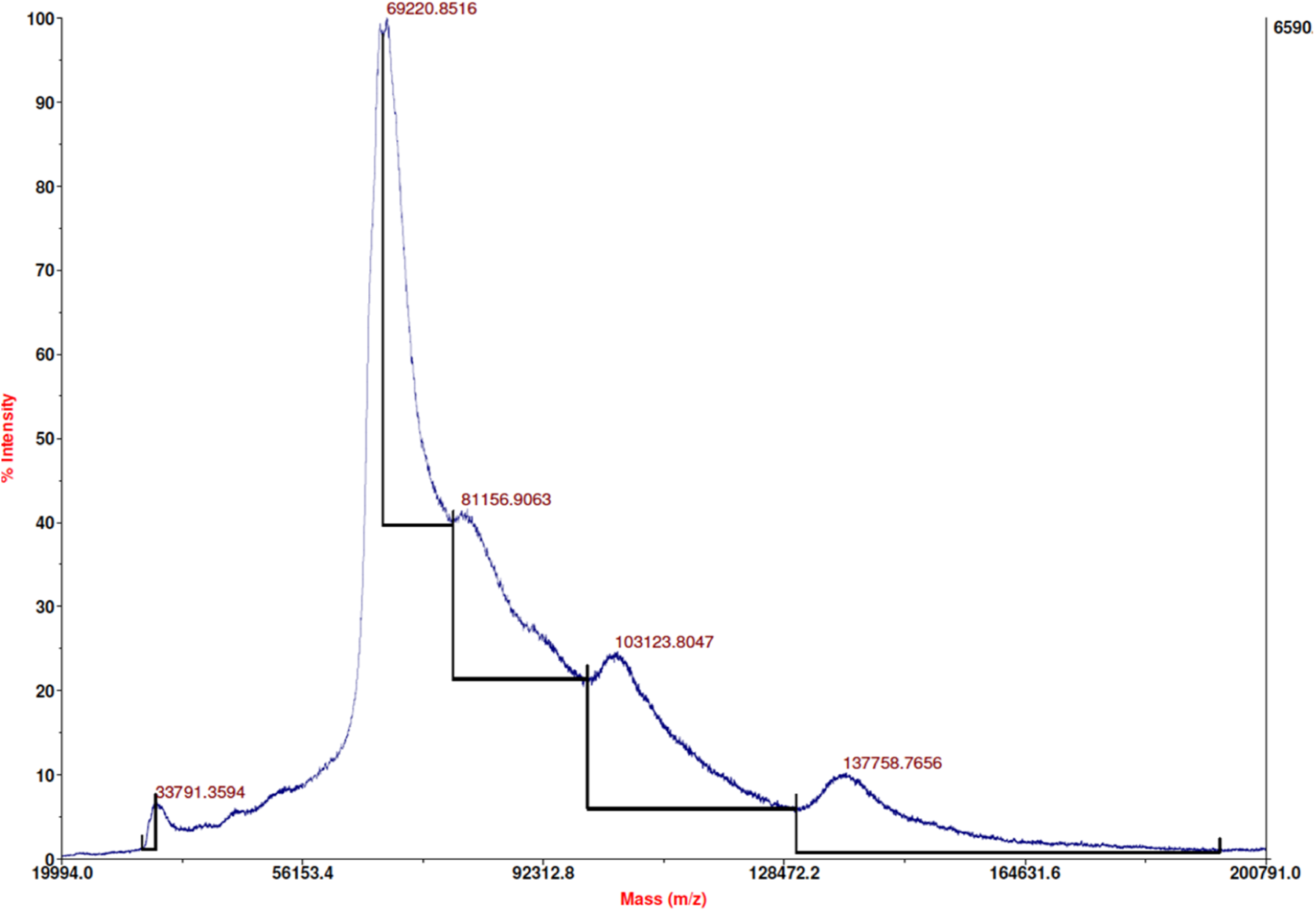
MALDI-TOF-MS spectrum of HSA-Tat/4T Complex.

**Figure S4.**
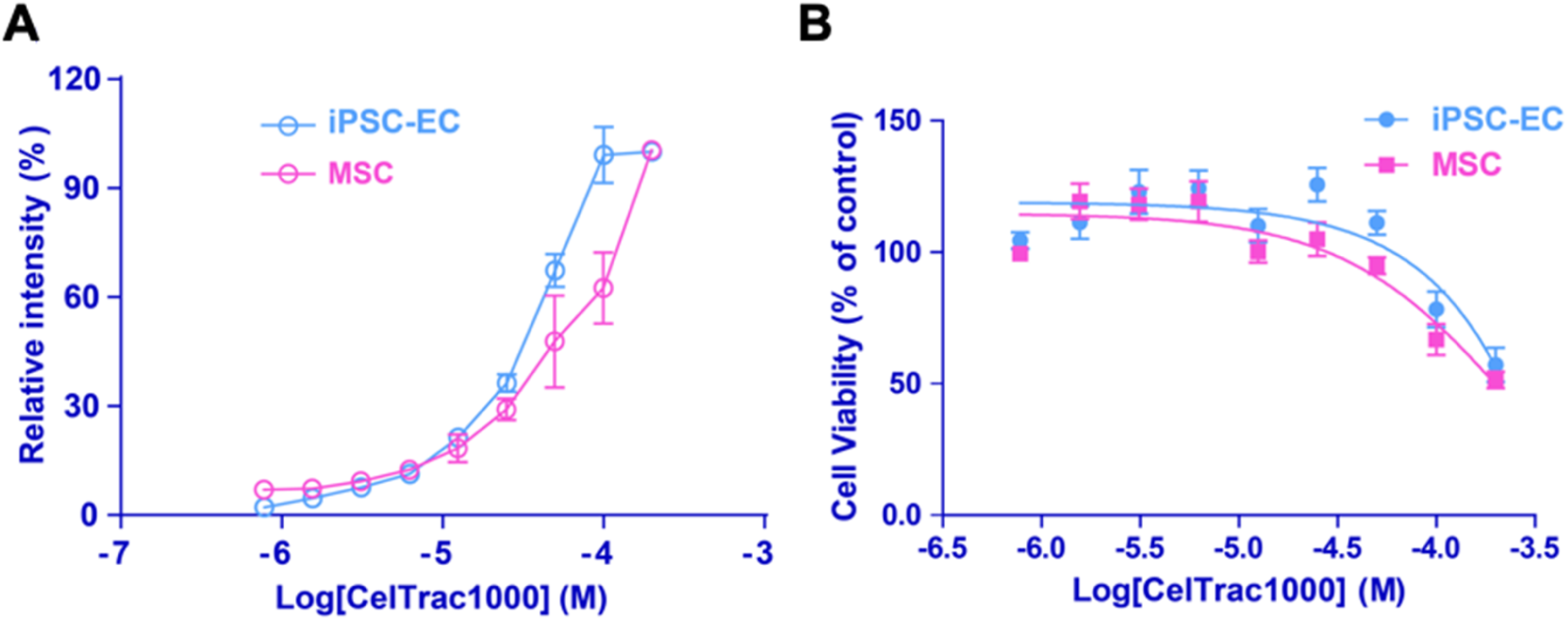
A) CelTrac1000 labeling concentration vs iPSC-ECs/MSCs relative NIR-II fluorescent intensity. B) CelTrac1000 iPSC-ECs/MSCs cytotoxicity analysis.

**Figure S5.**
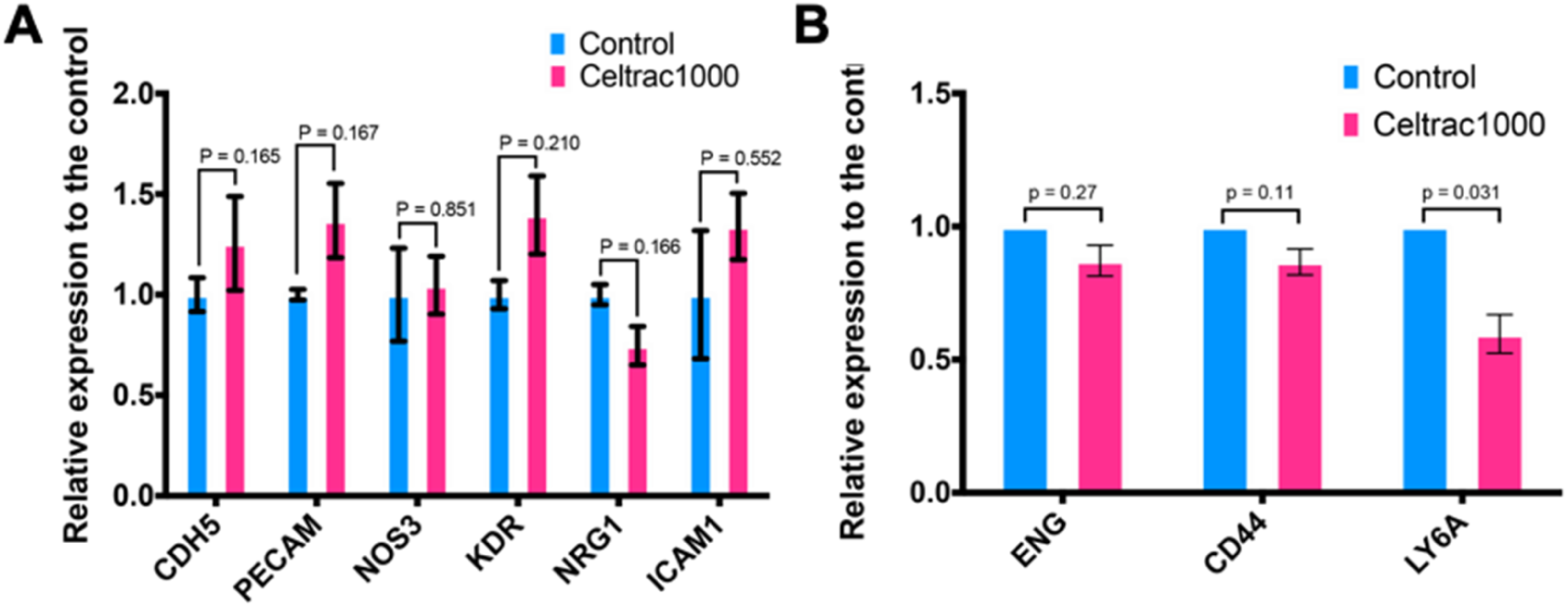
Comparison of gene expression between CelTrac1000 labeled iPSC-ECs (A) or MSC (B) and unlabeled control group. N = 3 per group.

**Figure S6.**
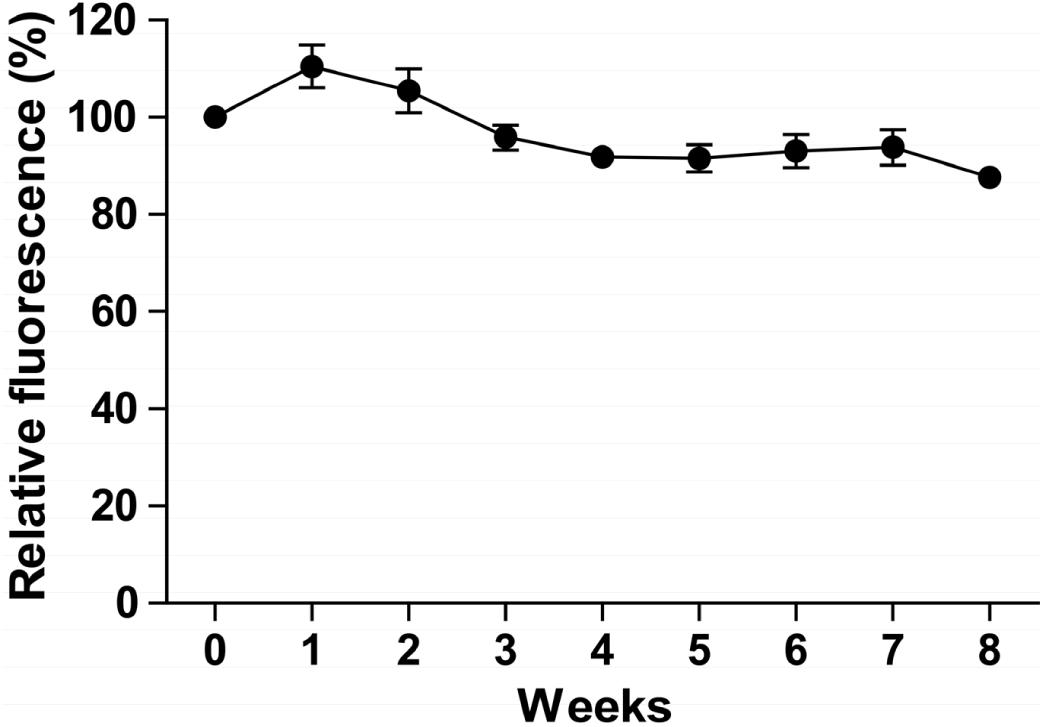
Plot of CelTrac1000 fluorescent changes in PBS buffer at 37 °C for 2 months.

**Figure S7.**
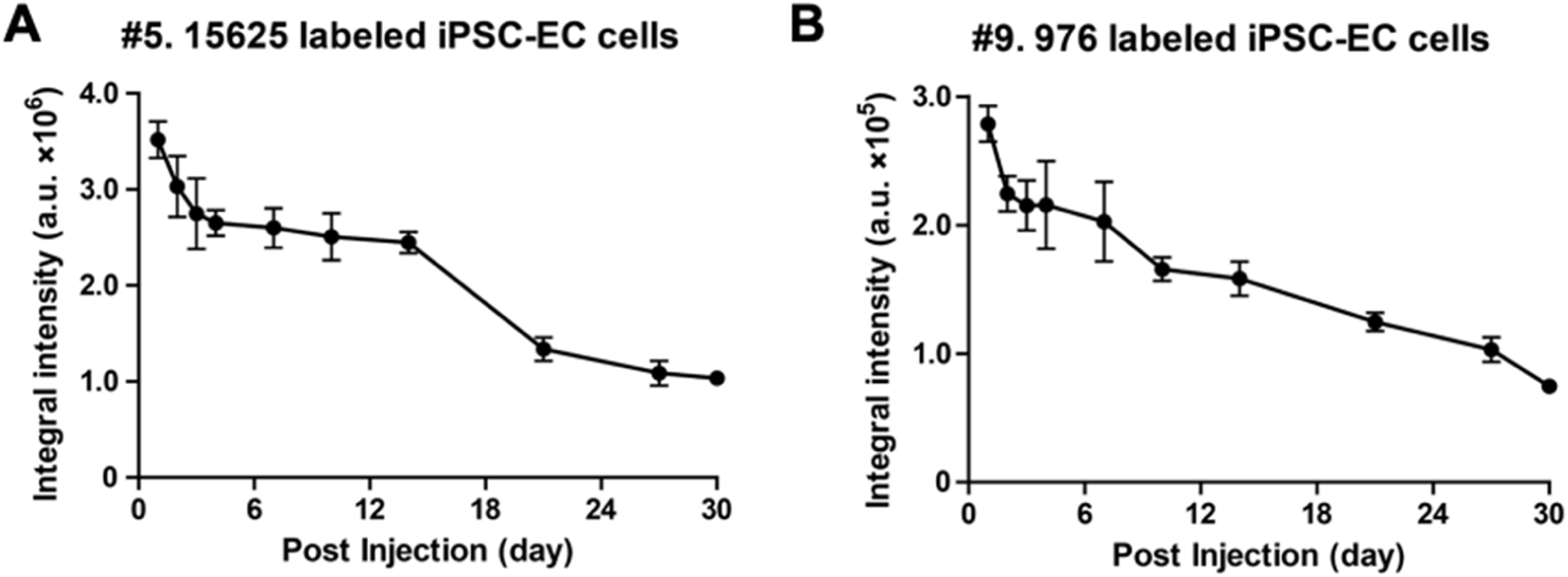
Plot of integral fluorescent intensities at spot 5 (A, 15,625 iPSC-ECs) and spot 9 (B, 976 iPSC-ECs) post injection.

## Appendix

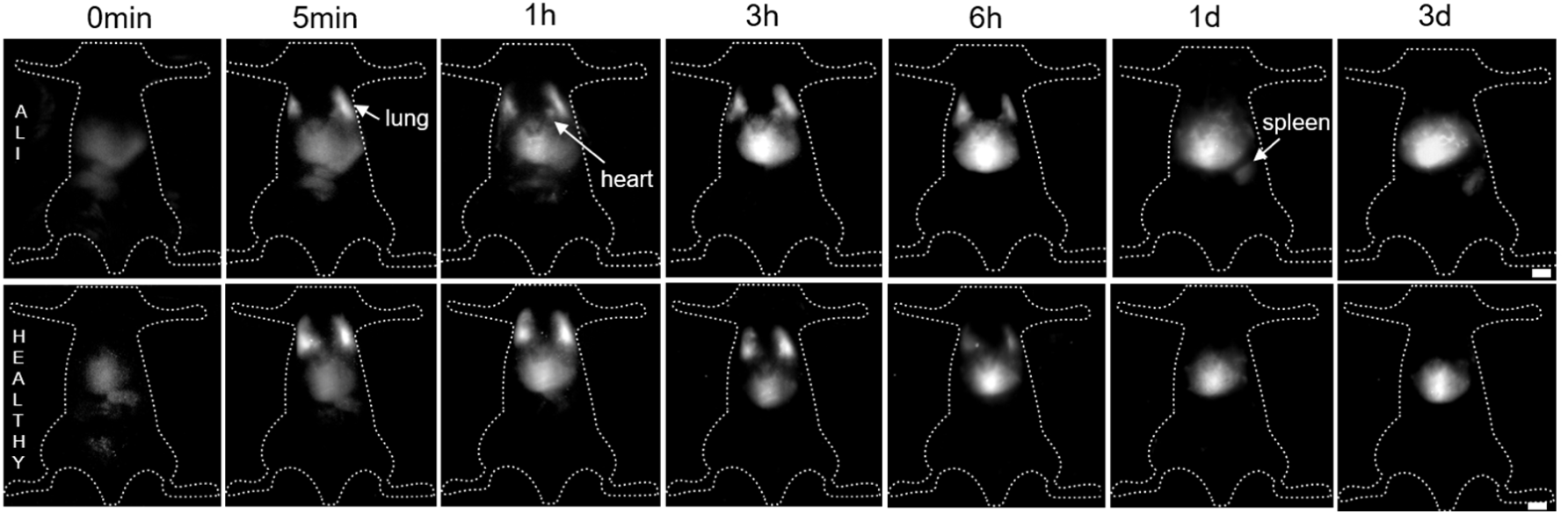

**1. The complete images of Fig. 4A**

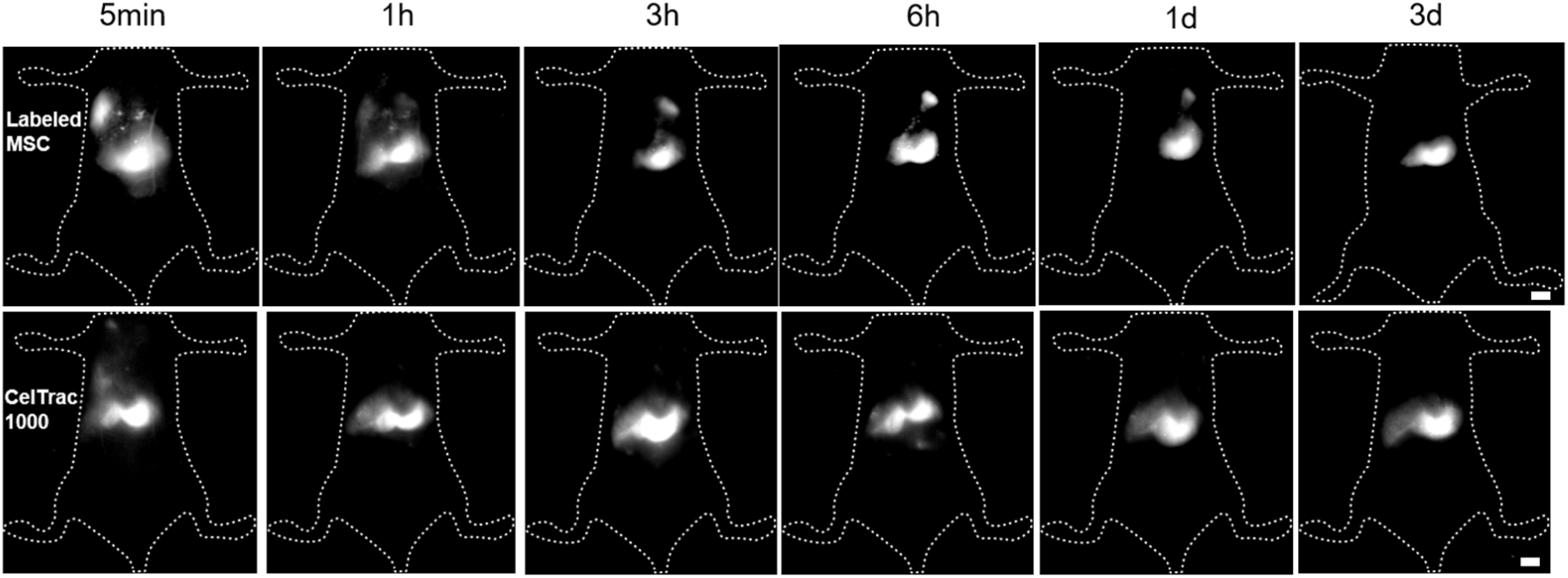

**2. The complete images of Fig. 5A**

### 3. The calculation formulas of cellular uptake and leakage of CelTrac1000

The cumulative amount of cellular uptake and leakage of CelTrac1000 MSCs and iPSC-ECs were calculated as follows:

#### Uptake

##### first 24h

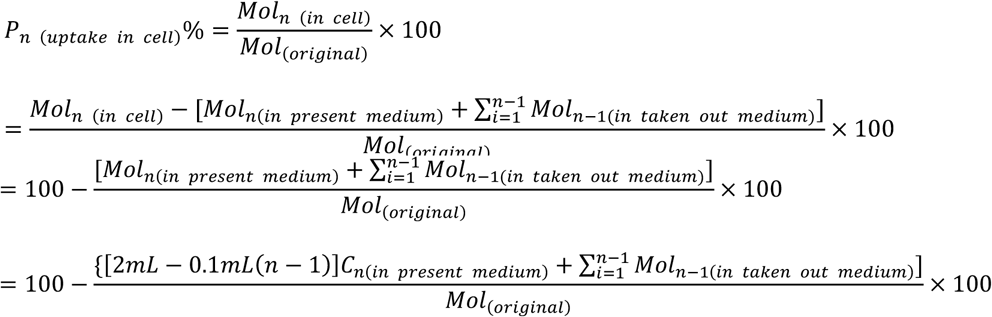

48h

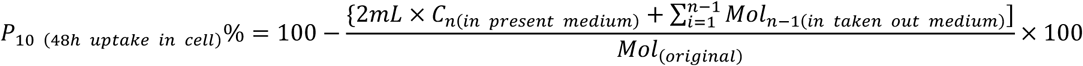

Where:

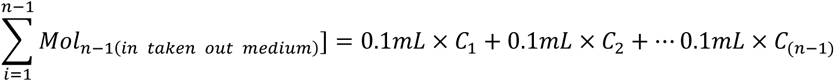

where *P* is the cumulative percentage of 4T/HSA-Tat in EC or MSC compare with the original, *Mol*_*n* (*in cell*)_ is the n^th^ mole amount of probe in cells, *Mol*_(*original*)_ is the original mole amount of probe which was put into mediums, *Mol*_*n(in present medium*)_ is the n^th^ probe (mole amount) in present medium, *Mol*_*n-1(in taken out medium*)_is the (n-1)^th^ mole amount of the probe in taken out medium. *C*_*n*_ is the probe concentration of the n^th^ taken out mediums (μmol/L).

#### Release

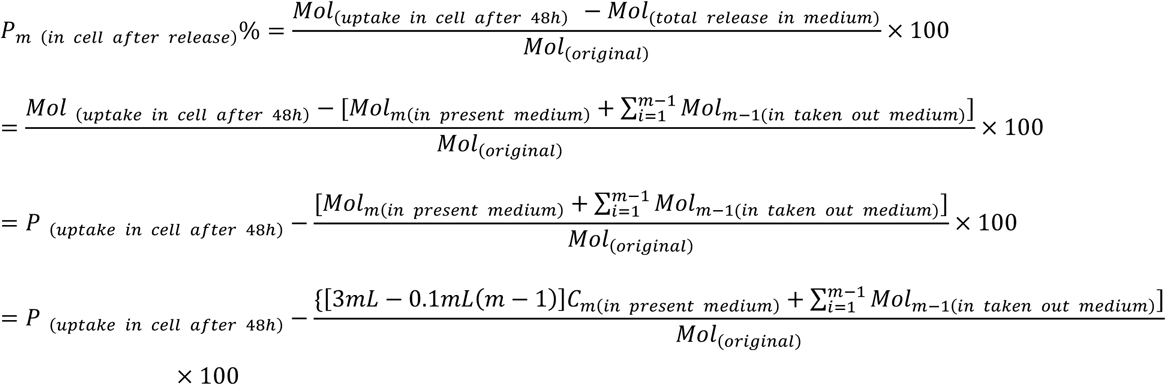

Where:

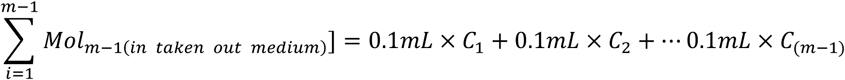

where *P* is the cumulative percentage of 4T/HSA-Tat in EC or MSC compare with the original, *Mol*_*m*(*uptake in cell after 48h*)_ is the m^th^ mole amount of probe in cells, *Mol*_(*original*)_ is the original mole amount of probe which was put into mediums, *Mol*_*m*(*in present medium*)_ is the m^th^ probe (mole amount) in present medium, *Mol*_*m-1*(*in taken out medium*)_is the (m-1)^th^ mole amount of the probe in taken out medium. *C*_*m*_ is the probe concentration of the m^th^ taken out mediums (μmol/L).

#### Release with IPS treated

##### First 6h

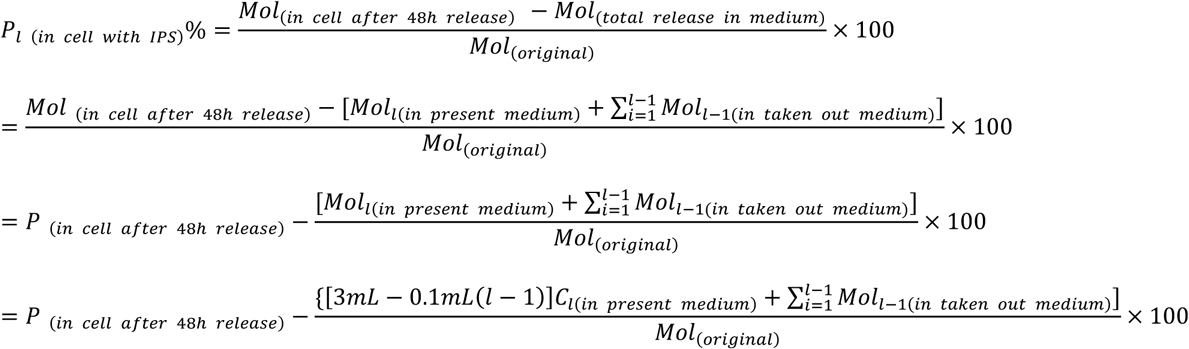

Where:

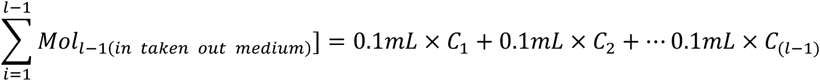

where *P* is the cumulative percentage of 4T/HSA-Tat in EC or MSC compare with the original, *Mol*_*l(in cell after 48h release*)_ is the l^th^ mole amount of probe in cells, *Mol*_(*original*)_ is the original mole amount of probe which was put into mediums, *Mol*_*l*(*in present medium*)_ is the l^th^ probe (mole amount) in present medium, *Mol*_*l-1*(*in taken out medium*)_is the (l-1)^th^ mole amount of the probe in taken out medium. *Cl* is the probe concentration of the l^th^ taken out mediums (μmol/L).

##### 6h to 48h

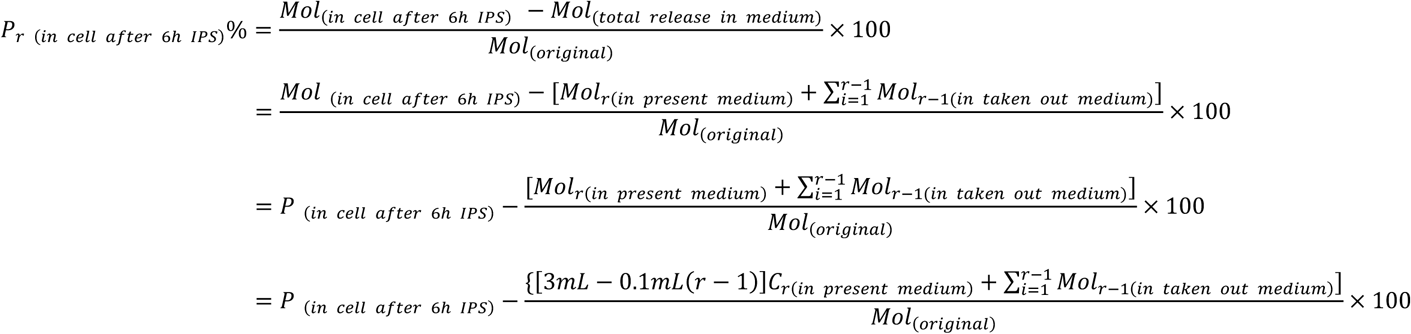

Where:

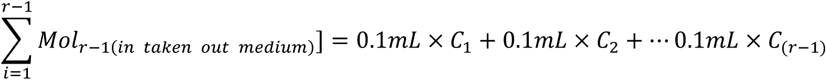

where *P* is the cumulative percentage of 4T/HSA-Tat in EC or MSC compare with the original, *Mol*_*r* (*in cell after 6h IPS*)_ is the r^th^ mole amount of probe in cells, *Mol*_(*original*)_ is the original mole amount of probe which was put into mediums, *Mol*_*r*(*in present medium*)_ is the r^th^ probe (mole amount) in present medium, *Mol*_*r-1*(*in taken out medium*)_is the (r-1)^th^ mole amount of the probe in taken out medium. *C*_*r*_ is the probe concentration of the r^th^ taken out mediums (μmol/L).

